# Near-atomic *in-situ* architecture and membrane-coupled dynamics of the *Vibrio cholerae* sheathed flagellum

**DOI:** 10.64898/2025.12.02.691832

**Authors:** Wangbiao Guo, Jian Yue, Jin Hwan Park, Diana Valverde Mendez, Jack M. Botting, Rajeev Kumar, Huaxin Yu, Avijay Sen, Jing Yan, Fitnat H. Yildiz, Jun Liu

## Abstract

The sheathed flagellum of *Vibrio cholerae* is a self-assembling membranous organelle that must coordinate axial assembly, sheath biogenesis, and rapid motor rotation. Here, we determine *in-situ* near-atomic structures of the sheathed flagellar motor inside intact cells. The motor anchors to the outer membrane through lipidated HL-rings without forming a membrane pore, thereby allowing axial assembly to drive sheath formation. Conserved LP-rings act as slide-rotary bushings that permit high-speed rotation within a dynamic envelope yet can constrict to seal the pore upon stress-induced ejection. We further show that stator activation requires a specific PomB-MotX interaction rather than peptidoglycan engagement. Together, these findings reveal how the distinctive architecture and dynamics of the sheathed flagellum promote *V. cholerae* motility, environmental survival, and persistent colonization of the human gut.

## Main Text

*Vibrio cholerae* is a highly motile, comma-shaped, Gram-negative bacterium that has evolved sophisticated mechanisms to navigate aquatic environments and colonize the human intestine, causing severe and highly transmissible diarrheal disease(*1–3*). Motility is central to its physiology and infection cycle, playing key roles in chemotaxis, colonization, biofilm formation, and virulence(*4–6*). Unlike model organisms such as *Escherichia coli and Salmonella enterica* that have unsheathed flagella, *V. cholerae* relies on a single polar flagellum encased in a membranous sheath and distinguished by *Vibrio*-specific features, including the H- and T-rings(*5*). These adaptations anchor the basal body (motor) to the outer membrane(*7–9*) and enable recruitment of sodium-driven stator complexes(*8, 10*) that generate exceptionally high torque(*11, 12*) and rapid rotation(*11–14*). However, how assembly of the polar flagellum is coordinated with sheath formation and how this specialized nanomachine operates within the dynamic bacterial envelope have remained poorly understood.

One critical factor in the survival of *V. cholerae* in aquatic habitats and persistent infection in human hosts is its ability to transition from a motile planktonic state to a sessile, surface-attached state(*15*). Recent studies demonstrated that under environmental stress such as nutrient depletion, cells actively shed the polar flagellum(*16*), leaving a flagellar outer membrane complex (FOMC) sealed by a plug-like structure(*9, 16–20*). However, the underlying molecular mechanism remains unclear.

To understand how the sheathed flagellum assembles, disassembles, and rotates within a dynamic and complex bacterial envelope, we integrated *in-situ* single-particle cryo-electron microscopy (cryo-EM) and cryo-electron tomography (cryo-ET) with genetic and biochemical approaches to capture conformational changes that coordinate flagellar assembly and disassembly in intact *V. cholerae* cells. We reveal how the sheathed flagellar motor anchors to the outer membrane via lipoprotein-based H- and L-rings, enabling formation of the membranous sheath in coordination with assembly of the flagellum. We also show that the sodium-driven stator complex exhibits remarkable flexibility, accommodating membrane fluctuations and specifically interacting with T-ring protein MotX for stator recruitment and torque generation. Finally, we propose that the motor achieves rapid rotation within the dynamic bacterial envelope through a novel slide-rotary bushing mechanism.

## Results

### Two distinct *in-situ* architectures of the flagellar motor in *V. cholerae*

To elucidate how the sheathed flagellar motor assembles and functions, we used an *in-situ* single-particle cryo-EM approach(*21*) to determine near-atomic resolution motor structures in intact *V. cholerae* cells **(Table S1)**. From 57,620 micrographs collected at the cell poles of a hyperflagellated Δ*flhG* mutant strain(*22*), we identified 155,760 flagellar motor particles through a combination of manual picking, deep-learning-based Topaz(*23*) particle selection, and iterative classification with cryoSPARC(*24*). The resulting *in-situ* structure at local resolutions of 2.7-3.8 Å reveals the MS-ring, rod, hook, and FOMC assembled around the rod. Notably, the FOMC is composed of the well-resolved L-, P-, H-, and T-rings **(Fig. S1-3)**. We also resolved the C-terminal domain of PomB (PomB_C_) at ∼5 Å resolution while bound to the T-ring. The rest of the stator is not resolved, likely due to its flexibility. This structure represents the fully assembled state of the flagellar motor **(Fig. 1a-c)**.

**Fig. 1:**
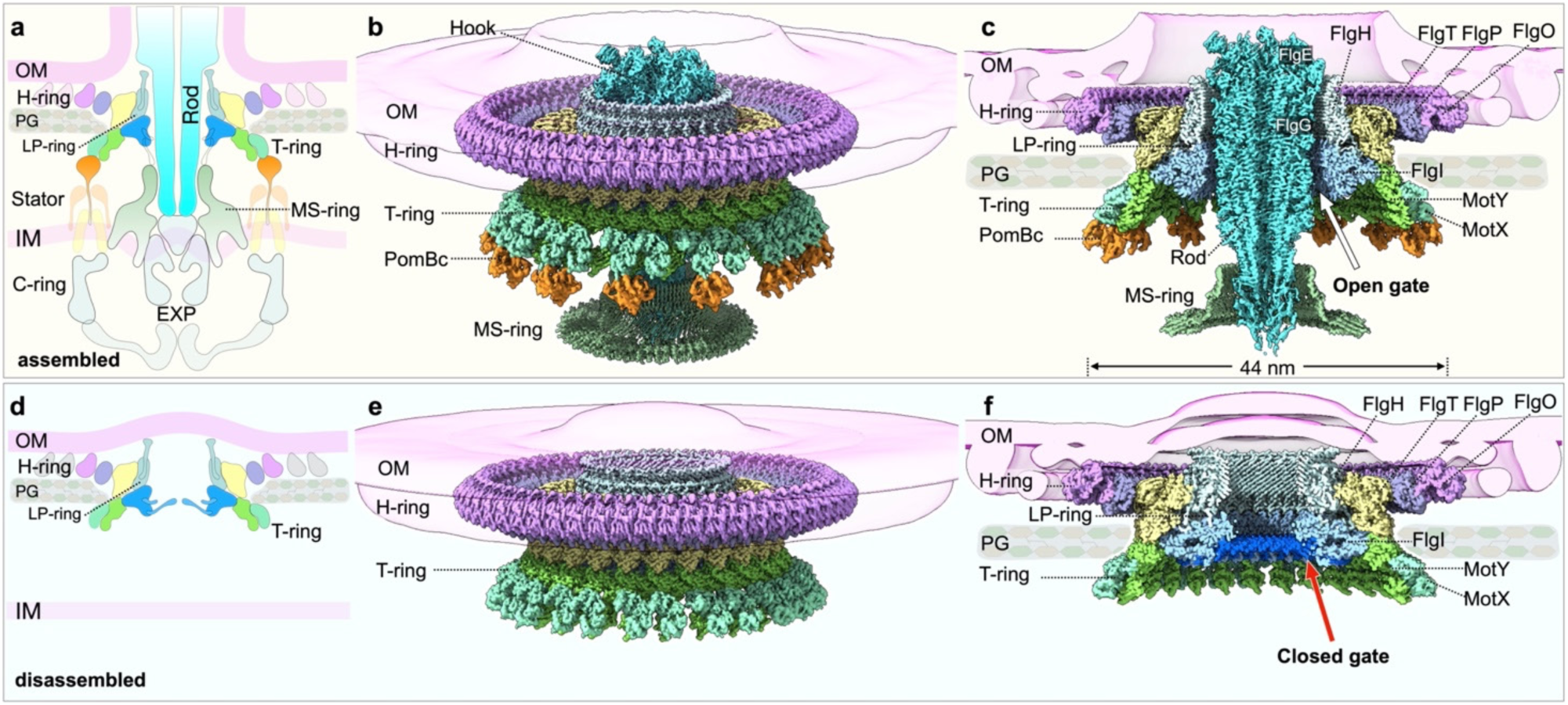
*In-situ* cryo-EM structures of the *Vibrio cholerae* flagellar motor. **(a)** Schematic model of the fully assembled flagellar motor in *V. cholerae*. **(b, c)** Overall (b) and cross-section (c) view of flagellar outer membrane complex (FOMC) at the assembled flagellar motor. The motor consists of the rod, hook, MS-ring, and FOMC, which includes the L-, P-, H-, and T-rings. The distal rod and its junction with the hook are formed by FlgG and FlgE, respectively. The proximal rod is surrounded by the periplasmic portion of the MS-ring (dark green). LP-rings are composed of FlgH (light cyan) and FlgI (light blue). The H-ring is assembled from FlgT (yellow), FlgP (dark purple), and FlgO (magenta), with additional unresolved peripheral densities. The T-ring consists of MotY (green) and MotX (light cyan), with the 13 stator PomB C-terminal dimers (orange) located below it. **(d)** Schematic model of the disassembled flagellar motor. **(e, f)** Overall view (e) and cross-section (f) of FOMC at the disassembled state. In this state, the central rod-hook complex is ejected, leaving a remnant of the FOMC. A closed gate is observed at the bottom of the P-ring (highlighted in blue), while a curved outer membrane cap covers the top of the FOMC. Other components including T-ring and H-ring are similar to those in the assembled motor but lack PomB_C_ binding.

We further determined a second, *in-situ* structure of the FOMC at near-atomic resolution after selecting 43,759 particles lacking the filament, hook, and rod. We term this orphan FOMC the closed state due to the gate-like density at the base of the P-ring and outer membrane closure at the top of the L-ring **(Fig. 1d-f)**. Given its similarity to the relic subcomplex(*9, 16–20*), we hypothesized that the closed FOMC represents a post-disassembly state. To test our hypothesis, we utilized live-cell fluorescence microscopy and cryo-ET to visualize wild-type *V. cholerae* under nutrient-rich and nutrient-depleted conditions. Nutrient depletion significantly increases the frequency of flagellar shedding and formation of the closed FOMCs **(Fig. S4)**, consistent with previous observations of flagellar ejection(*9, 16–20*). We therefore designate this closed FOMC structure the disassembled state.

These near-atomic *in-situ* structures enabled *de novo* modeling of the large assemblies comprised of more than 12 flagellar proteins and ∼340 subunits **(Fig. S5)** in two distinct functional states, providing a framework for understanding the molecular basis of assembly, disassembly, and function of sheathed flagella.

### The LP-rings undergo extensive remodeling that enables flagellar assembly and disassembly

In the assembled state, the L-ring is spatially separated from the outer membrane, which arches upward to form the membranous sheath that encloses the hook and filament **(Fig. 2a, b)**. In the disassembled state, the outer membrane bends slightly upward, sealing the top of the L-ring, where the hydrophilic hairpin loop of FlgH (residues 141-148) forms polar contacts with the inner leaflet **(Fig. 2e, f)**.

**Fig. 2:**
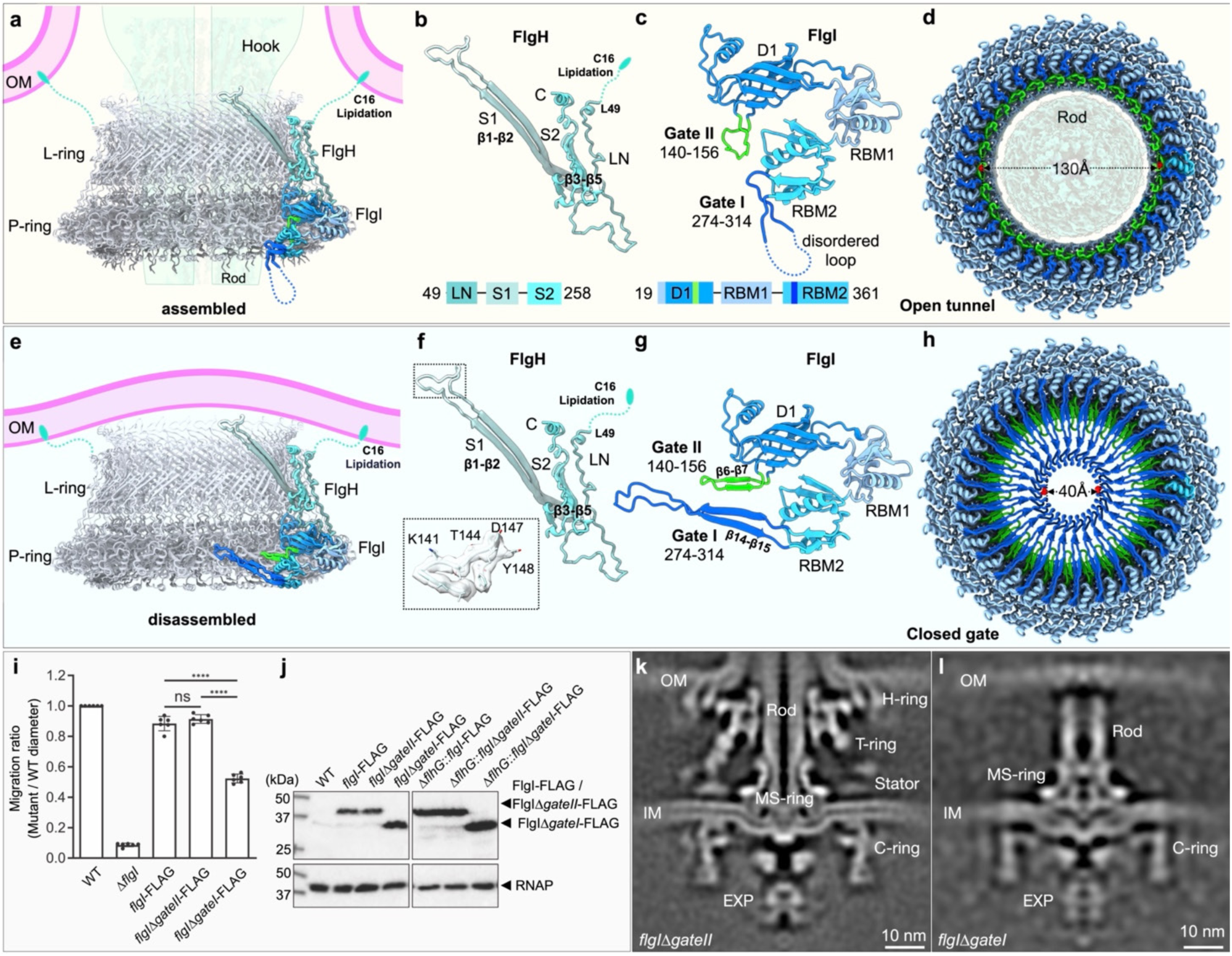
LP-rings during flagellar assembly and disassembly. **(a)** Structural model of the LP-rings in the assembled motor with one FlgH and one FlgI subunit highlighted. The unresolved flexible loop of FlgH (residues C16-G48, dashed line) contains the lipidated Cys16 that anchors the LP-rings to the OM. The rod-hook complex is shown as transparent cartoon. **(b)** FlgH monomer model, comprising an extended N-terminal loop (LN domain), an inner β-barrel (S1 domain), and an outer three-stranded β-barrel (S2 domain), along with N-terminal lipidation loop. **(c)** FlgI monomer consisting of an N-terminal D1 domain and two C-terminal ring-building motifs (RBM1 and RBM2). **(d)** Bottom view of LP-rings showing the ∼130 Å tunnel, forming an open tunnel for rod passage in assembled state. **(e)** Structural model of the LP-rings from the disassembled motor, where the overall structure is covered by a bent OM cap. **(f)** FlgH monomer from the disassembled motor, identical to that in the assembled state. Membrane-facing residues 141-148 are highlighted. **(g)** FlgI monomer from the disassembled state. Two gate-like loops are highlighted: GateI (residues 274-314, blue), and GateII (residues 140-156, green), in which GateI form a long antiparallel β-sheet at the base of the ring and Gate II forms a shorter β-sheet above it. **(h)** Bottom view of the disassembled LP-rings showing the central lumen narrowed to a closed gate of ∼40 Å in diameter. **(i)** Soft agar motility assays of *flgI* gate mutants, in which GateI or GateII was replaced by a dipeptide glycine-serine (GS) linker. Migration zones from six replicates were quantified (**, *p* ≤ 0.0001; ns, not significant). **(j)** Western blot analysis of *flgI* mutants in wild-type and Δ*flhG* backgrounds, with all alleles C-terminally FLAG-tagged. **(k, l)** Subtomogram averages of *flgI*Δ*gateII* (k) and *flgI*Δ*gateI* (l) mutants. Deletion of GateII has little effect on flagellar motor assembly (k), whereas deletion of GateI disrupts assembly, which stalls at the rod stage (l).

Notably, interactions between the L-ring and outer membrane in sheathed flagella are markedly different from those in unsheathed flagella **(Fig. S6a-c)**. As an example, the *S. enterica* L-ring forms an outer membrane pore through which the unsheathed hook and filament sequentially assemble **(Fig. S6b)**(*25–27*). Both hydrophobic L-ring-membrane interactions and lipidation of lipoprotein FlgH at Cys22 enable stable insertion of the L-ring(*26, 27*). By contrast, *V. cholerae* FlgH possesses a long, flexible loop (residues 16-48), not clearly resolved in our structures, allowing insertion of the lipid-modified Cys16 into the membrane without forming a pore **(Fig. S6a)**. This distinct anchoring mode represents a critical adaptation for sheathed flagella in *V. cholerae*.

The P-ring adopts distinct open and closed conformations corresponding to the assembled and disassembled states, respectively **(Fig. 2c, g)**. Each FlgI monomer comprises an N-terminal D1 domain and two C-terminal ring-building motif (RBM) domains, RBM1 and RBM2, that together form three concentric subrings: upper D1, outer RBM1, and lower RBM2.

In the closed conformation, residues H274-K314 (GateI, in RBM2) form a long antiparallel β-sheet, while residues S140-N156 (GateII, in D1) form a shorter β-sheet above it. These elements oligomerize into a narrow, gate-like structure with an inner diameter of ∼40 Å in the P-ring lumen, further sealed by additional densities likely corresponding to peptidoglycan **(Fig. 2d; Fig. S1k, l)**. In the open conformation, the distal rod occupies the central lumen, GateI is partially disordered and bent downward to embrace the rod, and GateII forms a flexible loop inserted into a cleft in the P-ring **(Fig. S6d, e)**.

Comparison of the two conformations indicates that during the transition from the assembled to disassembled state, GateI and GateII undergo ∼90° upward movements accompanied by a loop-to-β-sheet transition **(Fig. S6f, g)**. We propose that these gates act as molecular switches regulating the P-ring lumen during flagellar disassembly. The observed conformational changes are consistent with the flagellar ejection model(*9, 16–20*), in which rod ejection from the MS- and LP-rings triggers P-ring closure, thus preventing leakage of cytoplasmic contents.

To further probe the functional roles of Gates I and II, we constructed Δ*gateI* and Δ*gateII* mutants in which these regions were replaced by dipeptide (GS) linkers in both wild-type and Δ*flhG* backgrounds **(Fig. 2i, j; SFig. 7a-c)**. FLAG-tagged *flgI* variants were used to quantify protein abundance and decouple expression from function. Western blotting confirmed comparable FlgI expression across strains **(Fig. 2j)**. Functionally, the Δ*gateI* mutant exhibited ∼50% reduced motility, whereas Δ*gateII* displayed motility similar to wild-type **(Fig. 2i)**. Cryo-ET analyses reveal that in the Δ*gateI* mutant (Δ*flhG* background), ∼45% of the motors stalled during rod assembly without forming the FOMC, ∼27% formed intact flagella, and ∼28% showed FOMC remnants after disassembly. By contrast, in the Δ*flhG* background alone, only 0.1% of the motors stalled, ∼88% formed intact flagella, and ∼12% displayed FOMC remnants **(Fig. 2k, l; Fig. S7d, e)**. Together, these results demonstrate that GateI and GateII of the P-ring function as conformational switches that coordinate flagellar assembly and disassembly. Given the high sequence conservation of FlgI across bacterial species **(Fig. S6h)**, this open-closed transition likely represents a conserved mechanism underlying flagellar assembly and disassembly.

### Architecture of the membrane-bound H-ring

The H-ring is essential for flagellar assembly and sheath biogenesis in *Vibrio* species(*7, 28–30*). It comprises at least three components: FlgT, FlgO, and FlgP. FlgT forms a C26-symmetric ring encircling the LP-rings, while FlgO and FlgP assemble with C58 symmetry into two concentric outer membrane-associated rings **(Fig. 3; Fig. S8)**. In addition to these well-defined features, two peripheral ring-like densities are visible at the outermost region of the H-ring **(Fig. 1c, f)**.

**Fig. 3:**
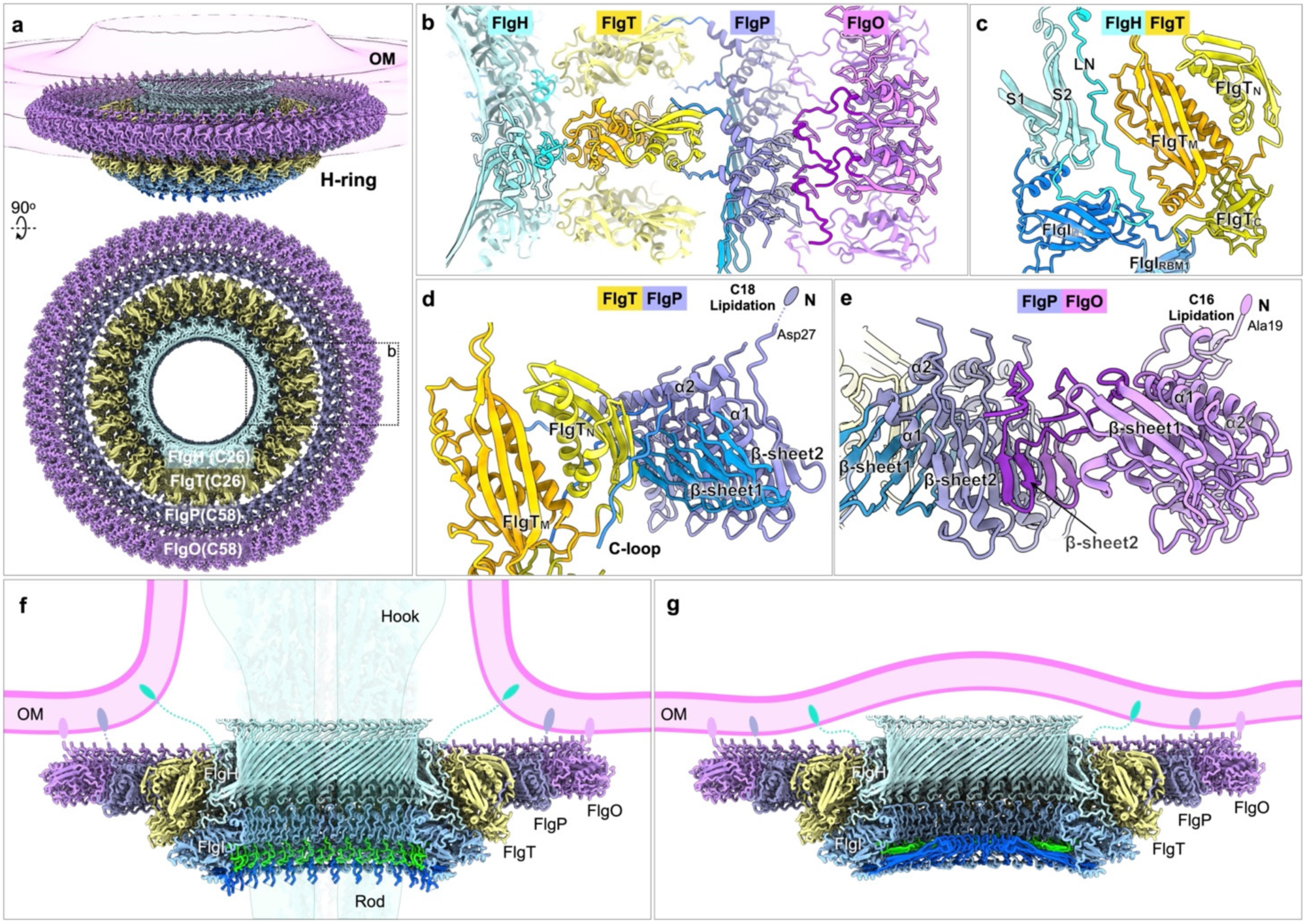
Architecture of the membrane-bound H-ring. **(a)** Tilt and top views of H-ring in the flagellar motor. The H-ring is composed of 26 copies of FlgT (yellow), 58 copies of FlgP (dark purple), and FlgO (magenta). **(b)** Zoomed-in top view of FlgH, FlgT, FlgP and FlgO. **(c)** Interaction between FlgH, FlgI and FlgT. **(d)** Interaction between FlgT and FlgP. The β-sheet1 in FlgP (blue) faces the FlgT_N_ with limited contact. The flexible C-terminal loop of FlgP (blue) extends along FlgT_N_, forming a more stable interaction. FlgP anchors to the OM via its N-terminal Cys18 lipidation (shown as a small ellipse dot). **(e)** Interface between FlgP and FlgO. The β-sheet2 (dark magenta) of FlgO engages β-sheet2 (dark purple) of FlgP, stabilizing the interface. FlgO anchors to the OM through its N-terminal Cys16 lipidation. **(f)** Cross-sectional view of the FOMC structure tethering to the OM (light magenta band) via the lipoproteins FlgH, FlgP, and FlgO in the assembled state. **(g)** Cross-sectional view of the FOMC structure tethering to the OM in the disassembled state.

FlgT is a 377-residue protein composed of three domains: FlgT_N_, FlgT_M_, and FlgT_C_ **(Fig. S8a, b)**. Compared with the crystal structure from *V. alginolyticus* (PDB: 3W1E)(*31*), the FlgT_N_ domain in *V. cholerae* is rotated upward, forming an additional concentric ring aligned with FlgT_M_ **(Fig. S8e)**. FlgT attaches to the LP-rings through interactions between FlgT_M_ and the LN loop of FlgH, burying ∼762 Å^2^ of the surface area per subunit **(Fig. 3b, c)**. The FlgT_C_ domain lies beneath FlgT_M_, acting as a scaffold for T-ring assembly **(Fig. S9)**.

FlgP adopts a compact sandwich-like fold consisting of two β-sheets (β1-β4-β5 and β2-β3) and two α-helices (α1, α2) **(Fig. S8a, c)**. The α1 helix is clamped between the β-sheets, with α2 positioned on top of the fold. β-sheet1 faces FlgT_N_ with minimal contact, while the C-terminal loop (residues 132-145) inserts into inter-subunit clefts within the FlgT ring, accommodating the symmetry mismatch between C58-symmetric FlgP and C26-symmetric FlgT **(Fig. 3b, d)**.

FlgO consists of two β-sheets (β1-β4, β5-β6) and two α-helices (α1, α2), with β-sheet1 interacting with both helices to form the core **(Fig. S8d)**. β-sheet2 extends toward FlgP and engages its β-sheet2, stabilizing the FlgO-FlgP interface **(Fig. 3d, e)**.

Both FlgO and FlgP are lipoproteins(*7, 29*) anchored to the outer membrane via conserved N-terminal lipidation at Cys16 (FlgO) and Cys18 (FlgP), as suggested by the diffuse densities observed near their N-termini in the outer membrane **(Fig. 3d, e; Fig. S8c, d)**. Together, these lipoprotein rings, FlgH, FlgO and FlgP, form a robust membrane-anchoring scaffold that secures the FOMC and couples it tightly to the outer membrane in both the assembled and disassembled states **(Fig. 3f, g)**. Given that mutants lacking the H-ring fail to assemble normal sheathed flagella and instead produce the non-motile periplasmic flagella(*28, 29*), the H-ring likely facilitates sheath formation by assisting the FOMC in stably anchoring to the outer membrane.

### Flagellar assembly is tightly coupled to sheath formation

To understand how the membrane-anchoring scaffold coordinates sheath formation with flagellar assembly, we performed cryo-ET analyses of polar sheathed flagella. Although long flagella were predominant, short, nascent flagella at different assembly stages were visible in both wild-type and Δ*flhG* cells. These shorter flagella were subjected to subtomogram analyses to capture early assembly intermediates.

The earliest stage of flagellar assembly visualized by cryo-ET shows the MS-ring and C-ring. Subsequently, formation of the rod provides a template for assembly of the open FOMC **(Fig. 4)**. As the hook elongates, it gradually pushes the outer membrane outward, forming a continuous sheath that envelops the growing hook **(Movie S1)**. Notably, the emerging hook extends perpendicularly from the outer membrane in a straight conformation rather than curved shape, suggesting that the straight conformation of the hook is energetically favorable in coordinating hook assembly and sheath formation. Interestingly, after flagellar ejection, the outer membrane completely seals the LH-rings, like the conformation before hook assembly **(Fig. 4c, d)**. Together, these early intermediates provide direct evidence that sheath biogenesis is intrinsically coupled to the concerted action of L-and H-ring formation and hook elongation.

**Fig. 4:**
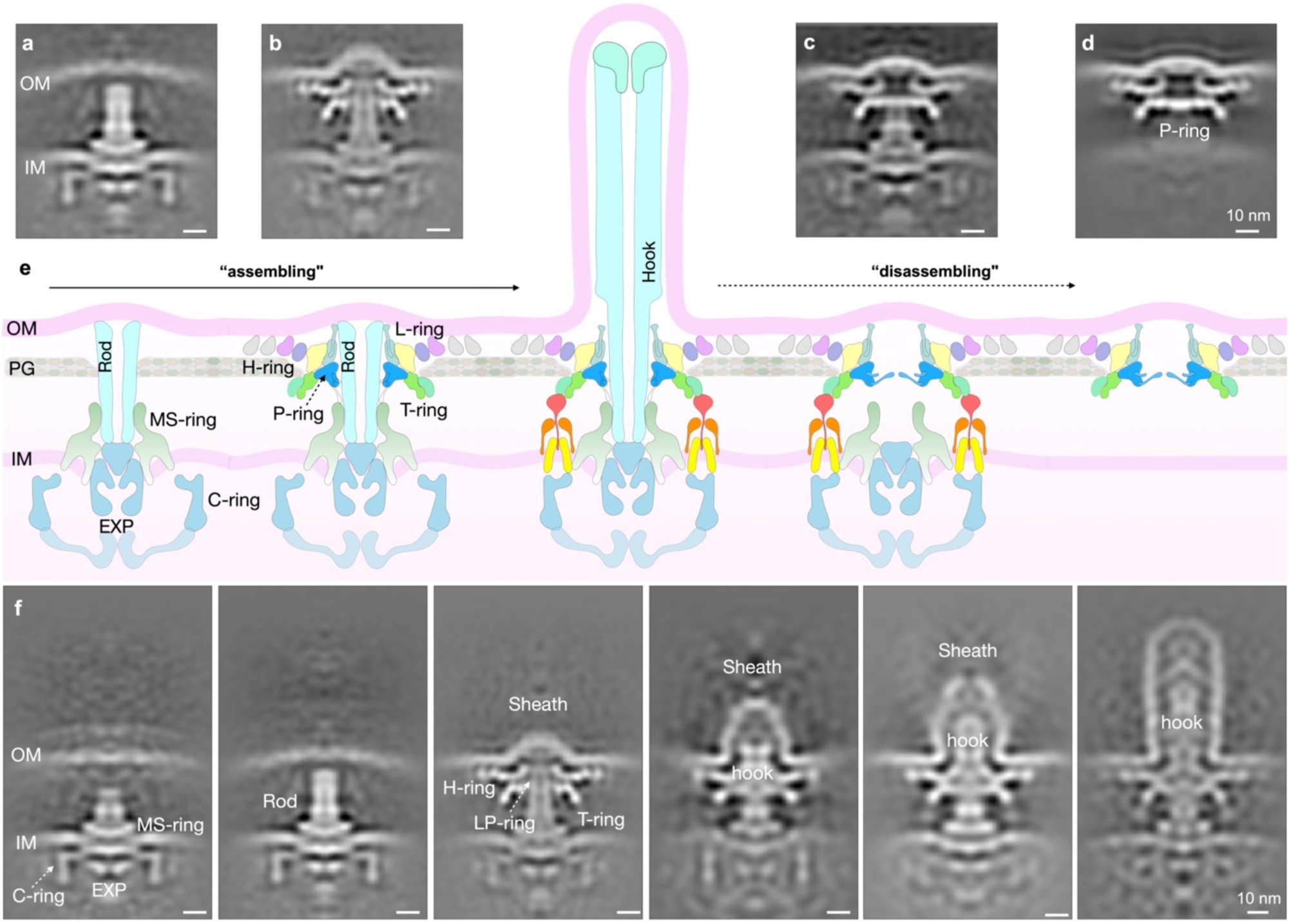
Key intermediates during flagellar assembly and disassembly. **(a, b)** Representative states of the flagellar motor assembly visualized by cryo-ET and subtomogram averaging. **(c, d)** Subtomogram averages of two representative states of flagellar motor disassembly. **(e)** Schematic model illustrating the simplified processes of flagellar assembly and disassembly. **(f)** Detailed view of the assembly intermediates. Flagellar motor assembly starts with the export apparatus (EXP), MS-ring, and C-ring, followed by rod assembly. The FOMC subsequently forms around the distal rod and anchors to the outer membrane (OM) via the LP-, H- and T-rings. As the hook elongates, it gradually pushes the OM outward, resulting in the formation of a continuous sheath that envelops the growing hook. The emerging hook extends perpendicularly from the OM in a straight conformation.

### Architecture of the T-ring and its interaction with the stator complex

The T-ring, composed of 26 MotX and 26 MotY subunits, resides beneath the LP- and H-rings and extends outward to contact the PomAB stator complex **(Fig. 5)**. Consistent with the crystal structure from *V. alginolyticus* (PDB: 2ZF8)(*32*), MotY consists of two domains: an N-terminal MotY_N_ and a C-terminal MotY_C_. MotY_N_ interacts with FlgT and FlgI, anchoring the T-ring to the basal motor **(Fig. S9a-c)**. Specifically, β3-β5 and α1-α2 of MotY_N_ face FlgT_C_, while a loop (residues 91-102) extends into the cleft between FlgT_C_ and FlgI-RBM1, forming hydrogen bonds and hydrophobic contacts. MotY_C_ adopts an α/β-sandwich fold containing a putative peptidoglycan-binding motif, although no peptidoglycan-like density is observed in our structure **(Fig. 5c)**.

**Fig. 5:**
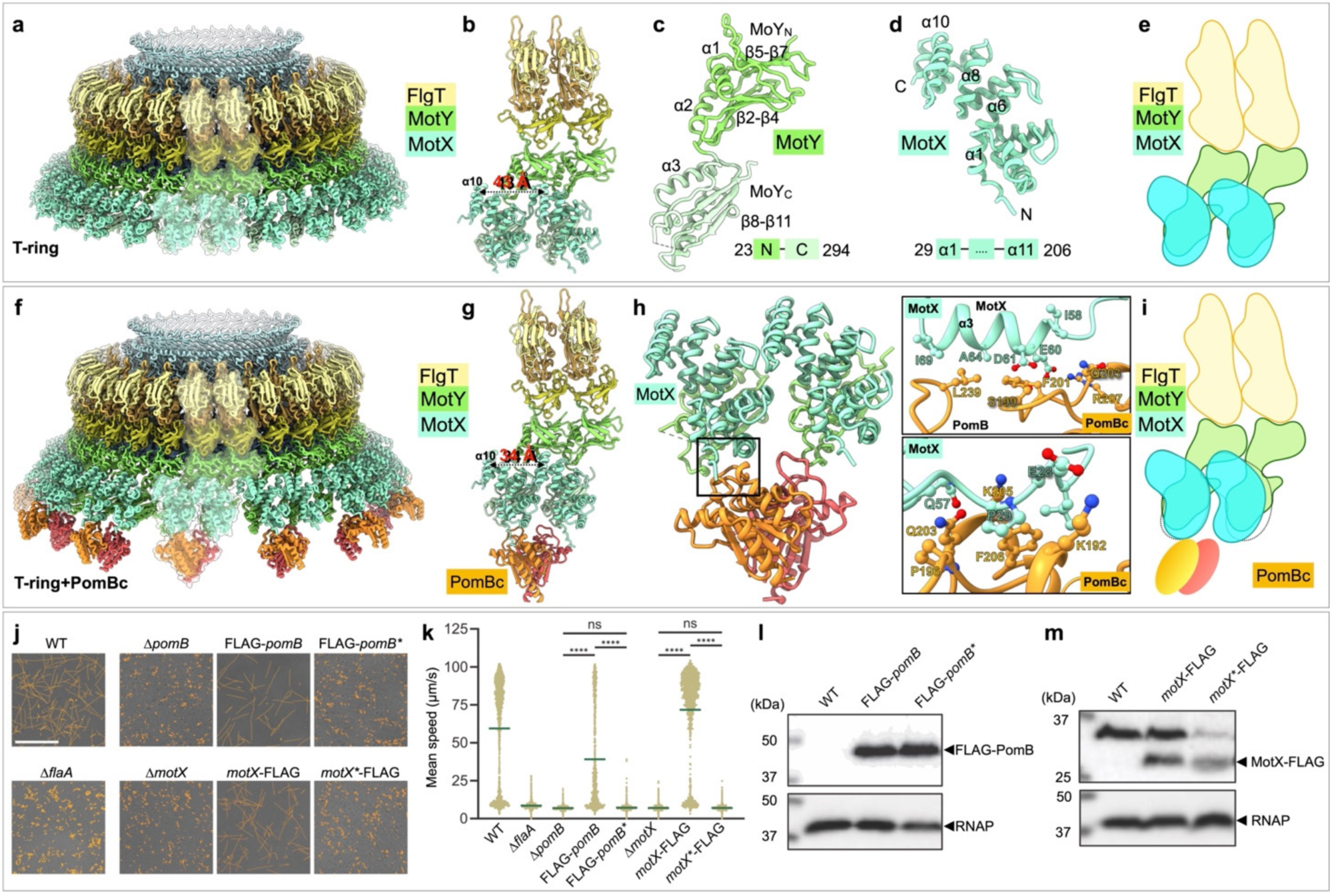
Architecture of the T-ring and its interaction with PomB. (**a**) T-ring structure without PomB_C_ binding. It is composed of 26 copies of MotY (green) and MotX (light cyan), sitting below FlgT and the LP-rings. (**b**) Zoom-in of two adjacent FlgT-MotY-MotX subunits, where the distance between the α10-helices of neighboring MotX subunits is 43 Å. (**c**) Monomer of MotY, consisting of N-terminal (MotY_N_) and C-terminal (MotY_C_) domains. (**d**) Monomer of MotX, adopting a helix-turn-helix fold comprising 11 α-helices. (**e**) Cartoon model of two FlgT-MotY-MotX complexes, illustrating the symmetrical assembly of adjacent MotX subunits. (**f**) T-ring structure with PomB_C_ binding, exhibiting a C13 symmetry. (**g**) Zoom-in of two adjacent FlgT-MotY-MotX-PomB_C_ subunits. PomB_C_ binding induces a rearrangement of MotX, reducing the α10-helix spacing to 34 Å. (**h**) Close-up of the MotX-PomB_C_ interaction. One PomB_C_ subunit (orange) interacts tightly with one MotX monomer, whereas the other (red) has limited contact. The key interaction sites are shown on the right. (**i**) Cartoon model of two FlgT-MotY-MotX-PomB_C_ subunits highlighting the rearrangement of MotX subunits upon PomB_C_ binding. (**j**) Representative snapshots of cell trajectories for strains with disrupted MotX-PomB interfaces obtained by single-cell tracking in liquid media. The *pomB* mutant (*pomB*^P196A,F201A,Q203A,F206A,L239A^) is denoted as *pomB^⁎^*, and the motX mutant (*MotX*^P29A,I58A,Q60A,D61A,A64R,I69A^) is denoted as *motX^⁎^*, scale bar = 50 μm. (**k**) Quantification of mean swimming speeds, where each dot represents an individual trajectory and the mean value is indicated by the dark green bar, adjusted *p* values ≤ 0.05 were considered significant, ****, *p* ≤ 0.0001, ns, not significant. (**l, m**) Western blot analysis of *pomB* (**l**) and *motX* (**m**) mutants, assessing protein expression and stability after mutation, with all *pomB* alleles tagged with an N-terminal FLAG epitope and all *motX* alleles tagged with a C-terminal FLAG epitope.

MotX, composed of tetratricopeptide repeat (TPR) motifs, adopts an 11-helix helix-turn-helix architecture **(Fig. 5d)**. Unlike MotY, MotX exhibits two conformations depending on its interaction with PomB **(Fig. 5e, i)**. In the absence of PomB_C_, the MotX-MotY complex maintains a symmetric C26 arrangement. Upon PomB_C_ binding, however, adjacent MotX subunits form dimers that engage PomB_C_ **(Fig. S9d, e)**. The inter-subunit distance between paired MotX monomers decreases from 43 Å to 34 Å. One MotX subunit forms a tight interface with the first PomB_C_ subunit of a PomB dimer, involving the N-terminal loop (residues 25-31) and α3 helix (residues 58-69) of MotX and loops 194-208 and 238-240 of PomB_C_. The second PomB_C_ monomer interacts weakly, inserting into the cleft between adjacent MotX-MotY subcomplexes **(Fig. 5f-h)**.

To assess the functional importance of the MotX-PomB interactions, we generated point mutations in each protein in residues contributing to the MotX/PomB interface: MotX* (P29A, I58A, Q60A, D61A, A64R, I69A) and PomB* (P196A, F201A, Q203A, F206A, L239A) **(Fig. 5j-m; Fig. S10)**. The PomB* mutant was expressed at levels comparable to wild type but exhibits a non-motile phenotype with no swimming activity in soft agar. Similarly, the MotX* mutant also displays a non-motile phenotype, though with protein levels reduced to ∼50% of wild type. Given that PomB and MotX are core components required for torque generation by the *Vibrio* sheathed flagellar motor(*33*), we next examined how perturbing the MotX-PomB_C_ interface affects swimming performance. Using single-cell tracking, we quantified the distribution of trajectory-averaged swimming speeds. Wild-type cells exhibited pronounced heterogeneity with a bimodal profile consisting of a low-speed fraction (∼0-10 µm/s) and high-speed fraction (∼50-100 µm/s) **(Fig. 5k)**. The low-speed fraction overlapped the non-motile Δ*flaA* control and is best explained by passive, non-flagellar displacements consistent with Brownian motion under our imaging conditions(*34*). Δ*pomB* and Δ*motX* showed background-level displacements indistinguishable from those of Δ*flaA*, indicating that both proteins are essential for flagellar rotation **(Fig. 5j, k)**. Consistent with soft-agar migration phenotypes, FLAG-*pomB* displayed a modest reduction in swimming speed relative to wild-type, whereas FLAG-*pomB*\* was essentially non-motile and phenocopied Δ*pomB* and Δ*flaA*, indicating near-complete loss of flagellar propulsion. For MotX, *motX*-FLAG tracked similarly to wild-type, whereas *motX*\*-FLAG behaved like Δ*motX* and Δ*flaA*. These results indicate that disruption of MotX-PomB_C_ interaction severely impairs flagellar function and bacterial motility.

Together, these studies demonstrate that MotX-PomB interaction is essential for stator recruitment and flagellar rotation, providing the first direct evidence that MotX, rather than peptidoglycan, plays a central role in stator activation and torque generation in *V. cholerae*.

### Slide-rotary bushing mechanism

Powered by 13 sodium-driven PomAB stator complexes, the sheathed flagellar motor in *Vibrio* generates exceptionally high torque(*11, 12*) and rapid rotation(*11–14*) while adapting to the dynamic bacterial envelope. To probe these dynamics in the fully assembled state of the motor, we performed 3D classification and focused refinement to determine *in-situ* structures of the distal rod and hook at 3.4 Å resolution, enabling model building of FlgG and FlgE **(Fig. 6; Fig. S11)**. Further focused classification on the FOMC revealed distinct orientations and axial movements up to 20Å, indicating that the rod undergoes rotation as well as axial sliding relative to the FOMC **(Fig. 6a, b; Movie S2)**. Notably, the PomB_C_ dimer remains attached to MotX of the T-ring **(Fig. S11; Movie S2)**, suggesting that the PomAB stator complex possesses intrinsic flexibility that enables specific interaction between PomB_C_ and MotX. This model is consistent with previous findings that both PomB and its homolog MotB contain a flexible linker to function as an effective anchor(*35, 36*).

**Fig. 6:**
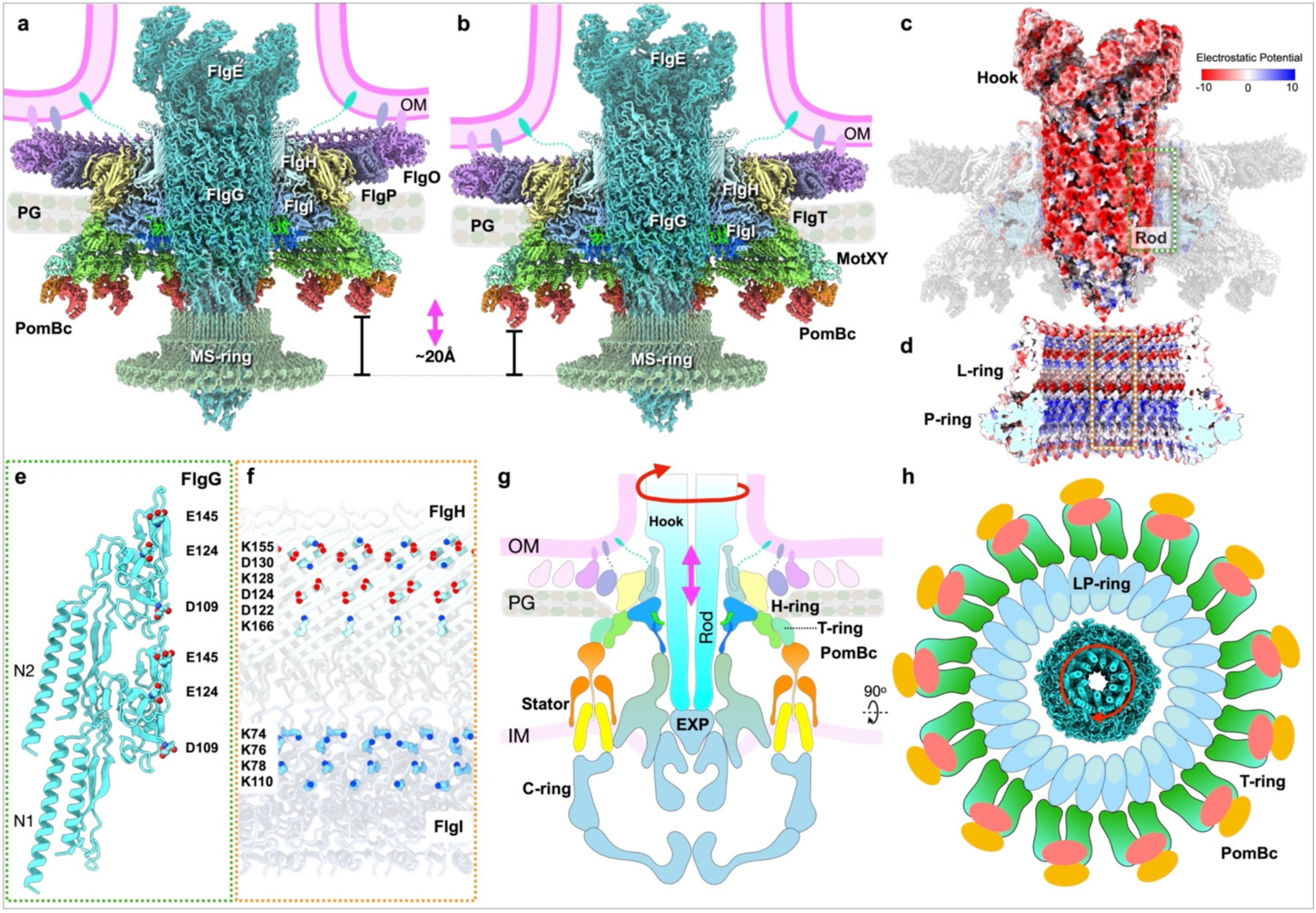
Distinct architectures of flagellar motor supporting a sliding-rotating model. **(a, b)** Overall structural models of the flagellar motor, including the rod (FlgG), hook (FlgE), FOMC, MS-ring, and the stator unit PomB_C_. 3D classification revealed multiple axial positions of the rod and FOMC, as reflected in the varying distance between the MS-ring and the FOMC. The two most divergent conformations differ by up to ∼20 Å, accompanied by synchronous fluctuations of the outer membrane. **(c)** Electrostatic analysis shows that the distal rod exhibits a fully negatively charged electrostatic surface potential. **(d)** The inner periphery of the LP-rings displays alternating negative and positive charged. **(e)** Key negatively charged residues mapped on the surface of the rod subunit FlgG. **(f)** Key positively charged residues located on the inner periphery of the LP-rings, contributed by FlgH and FlgI. **(g, h)** The sliding-rotating model of the flagellar motor from side view **(g)** and top view **(h)**.

The distal rod surface is enriched in acidic residues, generating a strongly negative electrostatic potential, whereas the inner wall of the LP-rings alternates between negatively and positively charged patches **(Fig. 6c, d)**. The P-ring forms the closest contact with the rod, and four lysine residues in the D1 domain of FlgI generate a ∼20 Å-wide, positively charged tunnel surrounding the rod **(Fig. 6e, f)**. These complementary electrostatic forces balance attraction and repulsion to maintain the coaxial alignment of the rod and stabilize its rotation, as proposed in *S. enterica*(*26, 27*). Moreover, the smooth, uniform surface of the distal rod allows the LP-rings to slide along the distal rod for ∼20 Å, effectively accommodating intrinsic fluctuations of the dynamic bacterial envelope. Together, these findings define a novel slide-rotary bushing mechanism by which the flagellar motor achieves rapid rotation within the dynamic bacterial envelope **(Fig. 6g, h; Movie S2)**.

## Discussion

The sheathed flagellum is a bacterial membranous organelle first discovered more than 70 years ago. Unlike the well-studied unsheathed flagella in *S. enterica* and *E. coli*, the sheathed flagellum in *V. cholerae* has evolved distinctive adaptations, including a membranous sheath, membrane-bound H-ring, and stator-associated T-ring, that enable high-speed and high-torque rotation as well as the dual sessile and motile lifestyles in *V. cholerae*. However, our understanding of the remarkable adaptations of the specialized membranous organelle has remained limited despite its critical roles in motility and pathogenesis. Moreover, the membranous sheath has long raised the questions of how a membrane can form around the flagellar filament and how it can accommodate rotation(*37, 38*). Integrating *in-situ* single-particle cryo-EM and cryo-ET with genetic and biochemical approaches, we now resolve the sheathed flagellum in unprecedented detail **(Fig. 1)**, revealing novel insights into how this complex membranous organelle assembles, rotates, and disassembles in intact *V. cholerae*. Our *in-situ* cryo-EM structures reveal that the L-ring in the *Vibrio* sheathed flagellum does not form a membrane pore but oligomerizes beneath the outer membrane at a distance maintained by a long, lipidated N-terminal loop of FlgH **(Fig. 3)**. Together with the large, lipidated H-ring, the motor firmly anchors to the inner leaflet of the outer membrane. These unique motor architectures and intermediates captured at the early stage of sheathed flagellar assembly support a model in which the growing hook pushes the outer membrane outward linearly, coupling sheath formation with hook assembly. The elongating filament subsequently continues to drive sheath extension **(Movie S1)**. Furthermore, given that the sheath is contiguous with the outer membrane and that both the H- and L- rings firmly anchor to the outer membrane, rotation of the filament, hook, and rod is likely independent from that of the sheath.

To generate high-torque and high-speed rotation, *Vibrio* species employ the T-ring to recruit and stabilize Na^+^-driven PomAB stator complexes(*14, 39*). In contrast to the prevailing model, in which the putative peptidoglycan-binding domain of PomB or its homolog MotB was thought to engage the cell wall(*40, 41*), our structures and point mutant experiments reveal that the putative peptidoglycan-binding domain does not bind peptidoglycan but rather primarily interacts with T-ring component MotX **(Fig. 4)**. It remains to be seen whether the stator complexes in other flagellar motors are also anchored to accessory proteins. MotB in *Campylobacterota* species, for instance, is enclosed within a cage-like structure and may be anchored to that structure rather than peptidoglycan(*42, 43*). Furthermore, our data provide evidence that the PomAB stator complex has intrinsic flexibility that maintains stable PomB-MotX interactions, ensuring robust stator recruitment and maximal torque generation **(Movie S2)**, consistent with the spring model previously proposed(*36, 44*).

To maintain high-speed rotation, the motor must function as a flexible transenvelope nanomachine to accommodate fluctuations of the dynamic bacterial envelope(*45*). We propose a slide-rotary bushing model to explain this complex and dynamic flagellar rotation **(Fig. 6; Fig. S11)**. Specifically, the membrane-bound LH-rings, composed of flexible lipoproteins, anchor to the outer membrane with variable curvature while sliding along the distal rod by up to ∼2 nm. This flexibility allows the motor to adapt to fluctuations of the dynamic envelope **(Movie S2)**. Similar sliding behavior has been observed at lower resolution in *V. alginolyticus*(*8*) and *Campylobacter jejuni*(*42*), suggesting a conserved strategy for maintaining high-speed rotation of sheathed and unsheathed flagella in complex and dynamic membrane environments.

It is well established that bacteria eject flagella as a strategy in adapting to changing environments or stress conditions(*46*). Recent studies have shown that FOMCs remain attached to the cell envelope after flagellar ejection in diverse bacterial species(*9, 16–20*). However, how the FOMC prevents leakage after removing the rod has remained unclear. Our *in-situ* structure reveals that, upon rod ejection, two flexible segments of FlgI (GateI and GateII) tilt upward to form β-hairpins that constrict the P-ring pore, thereby sealing the complex **(Fig. 2; Movie S2)**. Given the high conservation of FlgI across bacterial species **(Fig. S6h)**, this passive gating mechanism likely represents a general strategy for flagellar ejection that enables rapid transition from motile to sessile states.

Together, these results reveal the atomic architecture and dynamics of the *Vibrio* sheathed flagellum and establish a mechanistic framework for how this specialized membranous organelle assembles, rotates, and ejects, thus enabling high-speed rotation as well as the dual motile and sessile lifestyles of *V. cholerae*. These insights provide a structural foundation for understanding bacterial motility and pathogenesis and may guide strategies to inhibit flagellum-dependent infection in *V. cholerae* and related species.

## Acknowledgments

We thank Jennifer Aronson, Donghyun (Raphael) Park, and Shuaiqi (Phil) Guo for critical reading and editing of the manuscript. We thank Shenping Wu, Sarah Zhang, Jianfeng Lin, and Kangkang Song for helping on cryo-EM data collection. We thank Kai Zhang, Pengxin Chai, and Dongjie Zhu for advice and help on structure determination of the flagellar rod. We thank Siqi Zhu and Beile Gao for evolution analysis of the LP-rings and H-ring.

## Data availability

The atomic coordinates and corresponding density maps have been deposited in the Protein Data Bank (PDB) and the Electron Microscopy Data Bank (EMDB). The LP-rings in the closed state are deposited as PDB: 9YDT and EMD-72834, and in the open state as PDB: 9YDO and EMD-72815. Unless otherwise specified, all following components correspond to the open state. The H-ring subunit FlgT with C26 symmetry is deposited as PDB: 9YEF with EMD-72815. The H-ring subunits FlgO and FlgP with C58 symmetry are deposited as PDB: 9YDS and EMD-72833. The T-ring subunits MotX and MotY with C13 symmetry of MotX are deposited as PDB: 9YH0 and EMD-72947, while the C26 symmetry form is deposited as PDB: 9YEC and EMD-72847. The rod including FlgG, FlgF, and FlgE are deposited as PDB: 9YEE and EMD-72849. The RBM3 domain of FliF in the MS-ring is deposited as PDB: 9YDX and EMD-72840. The PomB_C_ dimer with MotX and MotY is deposited as PDB: 9YED and EMD-72848 **(Table S1)**. The overall FOMC in the closed state is deposited as PDB: 9YFG and EMD-72891, and in the open state with lower H-ring conformation as PDB: 9YH6 and EMD-72961. The open state with higher H-ring conformation is deposited as PDB: 9YH7 and EMD-72963.

## Funding

National Institutes of Health grants R01AI189907, R01AI087946 and R01AI132818 (W.G., J.Yue, R.K., and J.L.)

National Institutes of Health grants R01AI189907 and R37AI102584 (J.H.P. and F.H.Y.)

Alfred P. Sloan Foundation grant FG-2023-2085 (J.Yan and D.V.M)

National Institute of General Medical Sciences grant DP2GM146253 (J.Yan).

Burroughs Wellcome Fund Postdoctoral Enrichment Program (D.V.M.)

## Author contributions

Conceptualization: WG, JY, FHY, JL

Methodology: WG, JY, JHP, DVM

Investigation: WG, JY, JHP, DVM, RK, JMB, HY, AS

Visualization: WG, JY, JMB, JL

Funding acquisition: JY, FHY, JL

Project administration: JY, FHY, JL

Supervision: JY, FHY, JL

Writing – original draft: JY, WG, JHP

Writing – review & editing: JY, WG, JHP, FHY, JL

## Competing interests

Authors declare that they have no competing interests.

## Materials and Methods

### Strain construction

Plasmid construction was performed using Gibson Assembly (New England Biolabs, Ipswich, MA), with primers designed using the NEBuilder Assembly Tool and listed in **Table S2-3**. PCR amplification was performed using Q5 High-Fidelity DNA Polymerase (New England Biolabs) according to the manufacturer’s instructions. Plasmids for gene deletions were generated as previously described(*1*), while the knock-in constructs carrying point mutations or FLAG tags were synthesized and ordered from TWIST Bioscience (South San Francisco, CA). Deletion and knock-in plasmids were introduced into recipient strains by biparental mating using *E. coli* SM10 λpir or S17-1 λpir as the donor. Donor and recipient cultures were mixed at a 1:1 ratio and incubated on LB agar plates at 37°C for at least 6 hours. Transconjugants were selected on Luria-Bertani Miller (LB) agar containing rifampicin (100 μg/mL) and ampicillin (100 μg/mL). Successful gene deletions or knock-ins were confirmed by PCR using the gene-specific delA and delD primer sets.

### Bacterial culture

*V. cholerae* strains were cultured aerobically at 30°C in LB medium, composed of 1% tryptone, 0.5% yeast extract, and 1% NaCl. Overnight cultures were prepared by streaking bacterial strains from frozen glycerol stocks onto LB agar plates, followed by incubation at 30°C overnight. A single colony was then selected and streaked onto a fresh LB agar plate, which was incubated for an additional 12 hours.

### Cryo-EM sample preparation

*V. cholerae* cells were centrifuged at 2,000g for approximately 5 minutes in 1.5 mL tubes, and the resulting pellet was gently rinsed with phosphate-buffered saline (PBS). The pellet was resuspended in PBS to a final optical density (OD_600_) of 1.0 for plunge-freezing preparation. Cryo-EM samples were prepared using copper grids with a holey carbon support film (200 mesh, R2/1, Quantifoil) and Lacey C grids (200 mesh, Cu, Ted Pella, Inc.). The grids were glow-discharged for ∼30s prior to sample application. A 5 µL aliquot of the cell suspension was applied to each grid, which was then blotted with filter paper (Whatman^TM^) for approximately 6s. Grids were rapidly plunged into a liquid ethane-propane mixture using a Leica GP2 plunger. During the plunge-freezing process, the GP2 environmental chamber was maintained at 25°C and 90% relative humidity to optimize vitrification.

### *In-situ* cryo-EM data collection and processing

Cryo-EM specimen was imaged using a 300 kV Titan Krios electron microscope (Thermo Fisher Scientific) equipped with a field emission gun and a post-GIF K3 Summit direct electron detector (Gatan). Images were recorded in nanoprobe mode with the following settings: magnification at 81,000×, spot size at 5, illumination aperture at 1.22 µm, slit width at 20 eV, C2 aperture at 50 µm, and objective aperture at 100 µm. A customized multishot script was used to directly image the cell tips on the cryo-EM grids with SerialEM(*2*), applying defocus values from -1.6 µm to -2.2 µm. The total electron dose was ∼70 e^-^/Å^2^. The effective pixel size was 1.068 Å per physical pixel (super-resolution mode, half physical pixel size). A total of 57,620 micrographs were collected from Δ*flhG* cells.

All micrographs were initially motion-corrected(*3*) and then subjected to Patch CTF Estimation(*4*) in cryoSPARC **(Fig. S1)**. Approximately 18,000 motor particles were manually picked initially, followed by particle picking across all micrographs using the machine-learning-based algorithm Topaz(*5*). In total, 155,760 particles of the flagellar motor in the assembled, open state and 43,759 particles in the disassembled, close state were selected.

Micrographs were extracted with a box size of 1280 and binned to 320 in Fourier space. An initial model of the flagellar motor was generated by *ab initio* reconstruction using ∼5,000 manually picked particles, followed by heterogeneous refinement with C13 symmetry. Approximately 1,000 particles showing clear LP-ring density were selected as the initial model. After including all particles and performing homogeneous and local refinement with C13 symmetry, we obtained global flagellar motor structures in both assembled and disassembled states at bin 4, achieving a resolution of 8.68 Å (FSC 0.143), the resolution limit at this binning stage.

Subsequently, we focused on resolving the LP-ring structure **(Fig. S1f-l)**. Micrographs were further re-extracted with a box size of 448 pixels. To remove unnecessary junk particles, a focused mask was applied on LP-ring to conduct 3D classification. Local refinement was conducted using C26 symmetry, which yielded high-resolution reconstructions of the LP-ring at 2.69 Å for the assembled-state with 112,879 particles and 2.75 Å for the disassembled-state with 31,668 particles, serving as the structural foundation for subsequent refinement of the H-ring and T-ring.

To determine the symmetry of two additional ring-like densities, presumed to correspond to FlgP and FlgO components of the H-ring **(Fig. S2a-d)**. Micrographs were re-extracted with a box size of 720 pixels and binned to 512 because of the larger diameter of H-ring. We performed symmetry expansion of a prime number, e.g. C53, C57, followed by focused 3D classification on the H-ring. Among that, a C58-symmetric ring was identified, enabling us to assign C58 symmetry to both FlgP and FlgO. Further local refinement improved the resolution of FlgO and FlgP structures within the H-ring to 3.23 Å and 3.28 Å, respectively, in the assembled, open state and disassembled, close state. The H-ring component FlgT is positioned at the periphery of the LP-ring and exposes C26 symmetry. However, due to its connection with FlgP with C58 symmetry, the resolution of FlgT in the initial reconstruction was low. To improve this, symmetry expansion was performed using C29, based on the coordinates of FlgP and FlgO. Focused 3D classification on the FlgT/P region was then carried out. Particles that retained clear C26 symmetry for FlgT were selected and subjected to local refinement using C1 symmetry. This approach yielded focused reconstructions of the FlgT and FlgP regions at 3.75 Å resolution, in the disassembled, close state **(Fig. S2f, g)**. Compare the H-ring components (FlgT, FlgP and FlgO) at assembled and disassembled states, their conformations are identical at these two states, so we only showed one conformation. The H-ring at assembled state was further focused classified to evaluate its plasticity with OM **(Fig. S2h, j)**.

The T-ring is located below the H-ring and P-ring. To resolve its structure, we shifted the center to the T-ring region and re-extracted the particles using a box size of 720 pixels, binned to 512 pixels. Symmetry expansion was performed with C26, followed by focused 3D classification on the T-ring and PomB region. This classification separated the dataset into two classes: C13 particles with PomB density and C26 particles without PomB. Both subsets were subjected to local refinement focused on the T-ring and PomB region under C13 symmetry. This analysis yielded three distinct reconstructions: (1) 49% (53,628 of 109,959) of open-state particles exhibited PomB_C_ and C13-symmetric MotX, resulting in a 3.60 Å reconstruction; (2) 51% (56331 of 109,959) of open-state particles lacked PomB_C_ and displayed C26-symmetric MotX; (3) 25,553 particles of closed-state particles showed C13-symmetric MotX **(Fig. S2k-p)**.

The central rotating components of the flagellar motor were also resolved. The RBM3 domain of FliF in MS-ring is separated by 3D classification that focused on MS-ring region with C34-symmetry based on the coordinates of LP-ring particles. Local refinement of MS-ring region improved the resolution to 3.75 Å with 44,428 particles. Partial rod and hook region were solved with C1 symmetry. The LP-ring particles were symmetry expanded with C26 and then focus-classified on the rod region. One class exhibited prominent rod density. Subsequent rounds of local refinement allowed us to resolve the rod and hook structure of FlgG and FlgE at 3.44 Å resolution with 77,446 particles **(Fig. S3)**. The local resolution of the central helix region of the rod and hook is relatively reliable, whereas the peripheral region appears fuzzy, likely due to particle orientation bias. Signal-to-noise ratio iterative reconstruction method was further used to improve the distortion of our cryo-EM maps(*6*).

All cryo-EM maps were further processed using the deep learning-based refinement algorithm DeepEMhancer(*7*) to enhance visualization.

### Cryo-ET and subtomogram averaging

Frozen-hydrated specimens of Δ*flhG V. cholerae* cells were imaged on a 300 kV Titan Krios electron microscope (Thermo Fisher Scientific) equipped with a field emission gun and a K3 Summit direct electron detector (Gatan). Tilt series were acquired using SerialEM(*2*) in conjunction with the FastTomo script(*8*) and PaceTomo algorism(*9*), employing a dose-symmetric scheme. Imaging was conducted at a defocus of -4.8 µm, with a cumulative electron dose of ∼70 e^-^/Å^2^ distributed evenly across 33 tilts, ranging from -48° to +48° in 3° increments. Raw frames were corrected for beam-induced motion using MotionCorr2(*3*), followed by tilt alignment and stacking in IMOD(*10*). Tomographic reconstructions were generated from 8× binned tilt series using Tomo3D(*11*). Initial subtomogram extraction was performed on 8× binned tomograms, focusing on flagellar motors. Subsequent 3D alignment and classification were conducted using the i3 software package(*12, 13*) to refine particle coordinates and eliminate low-quality particles. Final subtomogram extraction was carried out on unbinned tomograms based on the refined positions, and further subtomogram averaging was conducted at bin4. A total of 112 tilt series were acquired for the *ΔflhGflgIΔgateI* mutant, 84 for the *ΔflhGflgIΔgateII* mutant, and 442 for the Δ*flhG* strain. A total of 273 tilt series were collected from wild-type cells under nutrient-rich condition (LB medium) and 81 tilt series for nutrient-deplete condition (M9 medium) **(Table S4)**.

### Modeling

The amino acid sequences of FlgH, FlgI, FlgT, FlgP, FlgO, MotY, MotX, PomB dimer, FlgG, FlgE and FliF were applied to AlphaFold3(*14*) to generate predicted structural monomer. The predicted models were fitted into the cryo-EM map using ChimeraX(*15*) followed by interactive manual adjustment in Coot(*16*) and real-space refinement in Phenix(*17*). Final models exhibited high stereochemical quality as assessed by Phenix validation tools **(Table S1)**. Sequence alignments were performed using Clustal Omega(*18*). ChimeraX(*15*) was used for cryo-EM map segmentation, visualization, and molecular surface rendering. Both ChimeraX(*15*) and Blender 4.4 were used for animation.

### Motility assays

Soft agar migration assays were conducted using LB medium containing 0.3% Bacto-Agar. Plates were prepared by pouring 150 mL of the medium into 150 × 15 mm petri dishes and left to dry at room temperature for a minimum of 24 hours before use. Overnight cultures grown on LB agar at 30°C were inoculated onto the motility plates by stabbing and incubated at 30°C for 16 hours. Migration zones were imaged using a ChemiDoc MP Imaging System (Bio-Rad, Hercules, CA), and migration diameters were normalized to wild type included on the same plate. Each experiment was performed with six biological replicates, and statistical analysis was carried out using one-way ANOVA followed by Tukey’s post hoc tests in GraphPad Prism (GraphPad Software, San Diego, CA).

Swimming behavior was assessed with slight modifications to a previously established method(*19*). Overnight cultures were grown in LB medium at 30°C for 16 hours. These cultures were then diluted 1:10 in fresh LB medium to obtain an appropriate cell density for imaging. A 50 µL aliquot of the diluted culture was placed on a concave microscope slide (Thermo Fisher Scientific) and covered with an 18 × 18 mm coverslip. Imaging was performed on an AxioImager Z2 Widefield Microscope (Carl Zeiss Microscopy GmbH, Jena, Germany) using a 20× objective lens. The focal plane was adjusted to approximately 50 µm below the coverslip, and time-lapse images were acquired at 54 frames per second over a 20-second interval. Cell trajectories were analyzed using the TrackMate plugin in ImageJ. Trajectories lasting between 0.2 and 2 seconds were included in the analysis, and mean swimming speed was calculated in µm/second. Each experiment was performed in triplicate (*n* = 3), with at least 200 trajectories analyzed per biological replicate. The data were analyzed using one-way ANOVA with Tukey’s correction for multiple comparisons in GraphPad Prism (GraphPad Software).

### Western blot

Whole cell lysates were prepared from the strains grown for16 hours in LB liquid medium. Cells were collected by centrifugation, and the pellets were resuspended in 2% (w/v) SDS, followed by heating at 95°C for 10 minutes. After centrifugation to remove debris, protein concentrations were determined using the BCA Protein Assay Kit (Pierce, Rockford, IL). Equal amounts of protein (40 µg) were loaded onto SDS-PAGE gels, separated by electrophoresis, and transferred to PVDF membranes using a semi-dry transfer apparatus (Bio-Rad). Membranes were blocked with 5% (w/v) nonfat dry milk and probed with a monoclonal anti-FLAG antibody (Sigma-Aldrich, St. Louis, MO). In parallel, a separate SDS-PAGE and transfer were performed under identical conditions and probed with an anti-RNAP monoclonal antibody (BioLegend, San Diego, CA) to assess equal loading. Protein bands were visualized by chemiluminescence using SuperSignal West Pico substrate (Pierce) and imaged with a ChemiDoc MP Imaging System (Bio-Rad). Band intensity was also quantified using the same imaging system.

### Flagella staining and fluorescence imaging

Flagella were stained using FM1-43FX membrane dye (Thermo Fisher Scientific) according to the manufacturer’s instructions with minor modifications. Overnight cultures grown in LB medium were harvested by centrifugation (2000g, 5 minutes) and resuspended in 5 µg/mL FM1-43FX. After 1 minute incubation on ice, formaldehyde was added to a final concentration of 4%, followed by an additional 10-minute incubation on ice. Cells were pelleted again (2000g, 5 minutes) and resuspended in HBSS (Hanks’ Balanced Salt Solution; 140 mM NaCl, 0.5 mM KCl, 0.03 mM Na_2_HPO_4_, 0.04 mM KH_2_PO_4_, 0.4 mM NaHCO_3_). Suspensions were mounted on 1% agarose pads and imaged using an AxioImager Z2 Widefield Microscope (Carl Zeiss Microscopy GmbH) with a 100× objective lens and the 38 HE eGFP filter set (excitation 450-490 nm, emission 500-550 nm) using an Axiocam 506 mono camera (Carl Zeiss Microscopy GmbH, 200 ms exposure time, depth of focus 0.90 μm).

### Cell labeling

Wild-type cells were imaged using confocal microscopy to observe the flagella shedding process. Cells from the frozen stock were grown shaking overnight at 30°C in LB medium (BD, Franklin Lakes, New Jersey). Cells were then back-diluted 1:100 in M9 media (Sigma-Aldrich, St. Louis, MO) supplemented with 0.5% Glucose and 0.25% Casamino acids (CAAs) (BD, Franklin Lakes, New Jersey), and grown shaking for 4-5 hours at 37°C. Cells (1, 2 and 3 µL) were pipetted onto the carbon side of gold EM-grids using cut pipette tips (to preserve the filaments). The grids were then inverted and placed on a glass bottom 96-well plate and topped with 100 µL of M9 media (no carbon source). FM1-43 Dye (*N*-(3-Triethylammoniumpropyl)-4-(4-(Dibutylamino) Styryl) Pyridinium Dibromide) (Thermo Fisher Scientific, Waltham, Massachusetts) was added to each well to a final concentration of 5 µg/mL and mixed well using a cut pipette tip.

### Confocal microscopy imaging

Fluorescence microscopy was performed using a Yokogawa W1 confocal scanner unit connected to a Nikon Ti2-E inverted microscope with a Perfect Focus System. Cells were excited by a laser at 488 nm with the corresponding filter. All fluorescent signals were recorded by a scientific complementary metal-oxide semiconductor camera (sCMOS, Hamamatsu). Confocal images were taken using a 100× silicone oil objective (Lambda S 100XC Sil numerical aperture = 1.35). Each field of view was 130 µm × 130 µm; 12-14 locations were imaged for each experiment as technical replicates. During time course, imaging was performed every 5 minutes, overnight (10-12 hours). Cells were incubated during imaging at 27°C using an incubation chamber. Minimal photobleaching was observed under specific imaging condition. All images presented are raw images rendered with Nikon NIS-Elements.

---

**Fig. S1:**
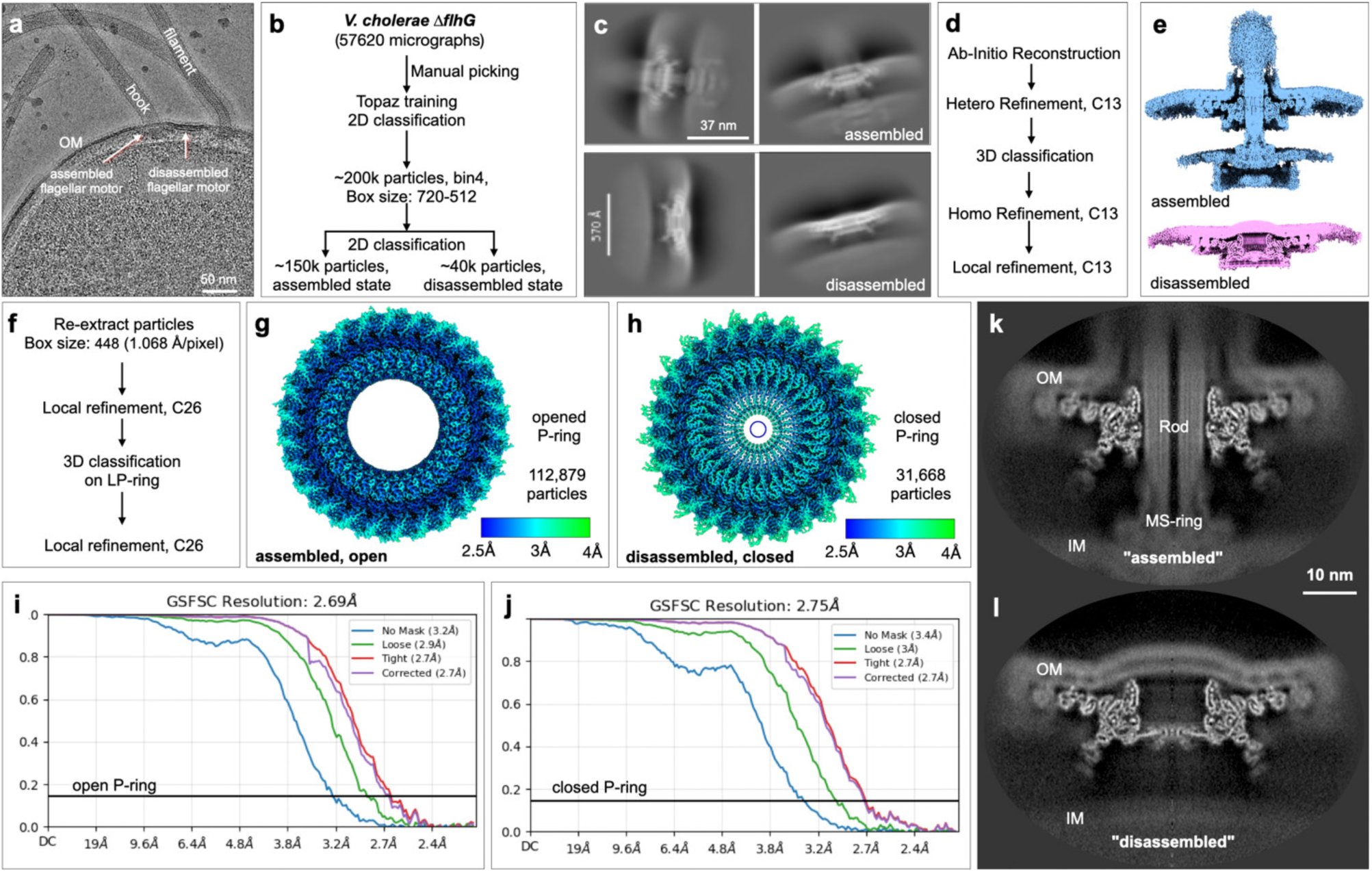
Workflow of *in-situ* structural determination of sheathed flagellar motor in *V. cholerae*. **(a)** Raw micrograph of a Δ*flhG V. cholerae* cell tip, showing flagellar motors in both fully assembled and disassembled states. Signals corresponding to the filament, hook, flagellar cap and flagellar motors are visible. **(b)** Workflow for *in-situ* cryo-EM data collection, particle picking, and 2D classification. **(c)** 2D classifications of assembled and disassembled flagellar motors at bin4. **(d)** Workflow for 3D reconstruction of the flagellar motor. **(e)** Cryo-EM maps of the global motor structure at bin4, where the complete flagellar motor components were visualized, including flagellar outer membrane complex (FOMC), inner membrane and C-ring. **(f)** Workflow for local refinement of the FOMC. **(g)** Local resolution map of the open LP-ring and FlgT in the assembled motor (112,879 particles, C26 symmetry). **(h)** Local resolution map of the closed LP-ring and FlgT in the disassembled motor (31,668 particles, C26 symmetry). **(i)** “Gold-standard” FSC curve (0.143 criterion) for the assembled motor with an open P-ring conformation. **(j)** “Gold-standard” FSC curve (0.143 criterion) for the disassembled motor with a closed P-ring conformation. **(k, l)** Cross-sectional views of the cryo-EM density maps of assembled and disassembled motor corresponding to panels (**g**) and (**h**).

**Fig. S2:**
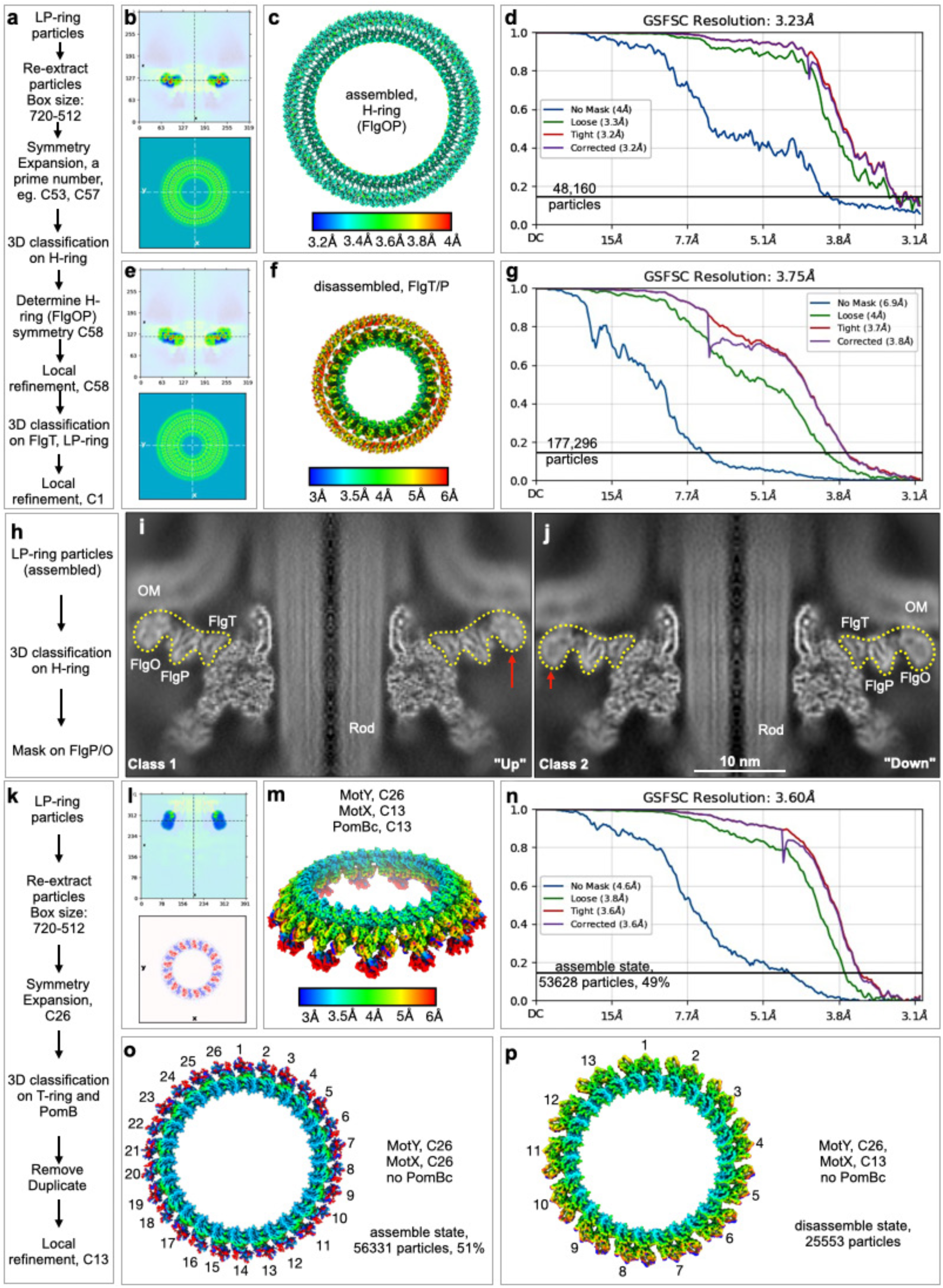
Data processing of the H-ring and T-ring. **(a)** Workflow for data processing and symmetry determination of the H-ring subunits FlgO and FlgP. **(b)** Focused mask on H-ring subunits FlgO and FlgP for 3D classification to determine their symmetry. **(c, d)** Local resolution maps and “gold-standard” FSC curves of FlgO and FlgP in assembled flagellar motors. The structures were determined with 48,160 particles. **(e-g)** Refining the structures of FlgT and FlgP with C1 symmetry to address symmetry mismatch. **(h-j)** Two distinct conformations of the H-ring and outer membrane reveal striking “up” and “down” fluctuations, while the LP-ring remains steady. **(k)** Workflow for structural determination of the T-ring and PomB. **(l)** Focused mask applied to the T-ring and PomB. **(m-n)** Local resolution map and FSC curve of the T-ring and PomB in the assembled flagellar motor. A conformation containing C13 MotX and C13 PomB dimers were sort out by 3D classification, including 53,628 particles (49% of the assembled dataset), reaching a global resolution of 3.6 Å. **(o)** 3D classification identified an assembled motor conformation lacking PomBC, but containing MotX with C26 symmetry. This class has totally 56,331 particles (51% of the assembled dataset). **(p)** Local resolution map and FSC curve of the T-ring and PomB in the disassembled motor. A conformation with C13 MotX with 25,553 particles.

**Fig. S3:**
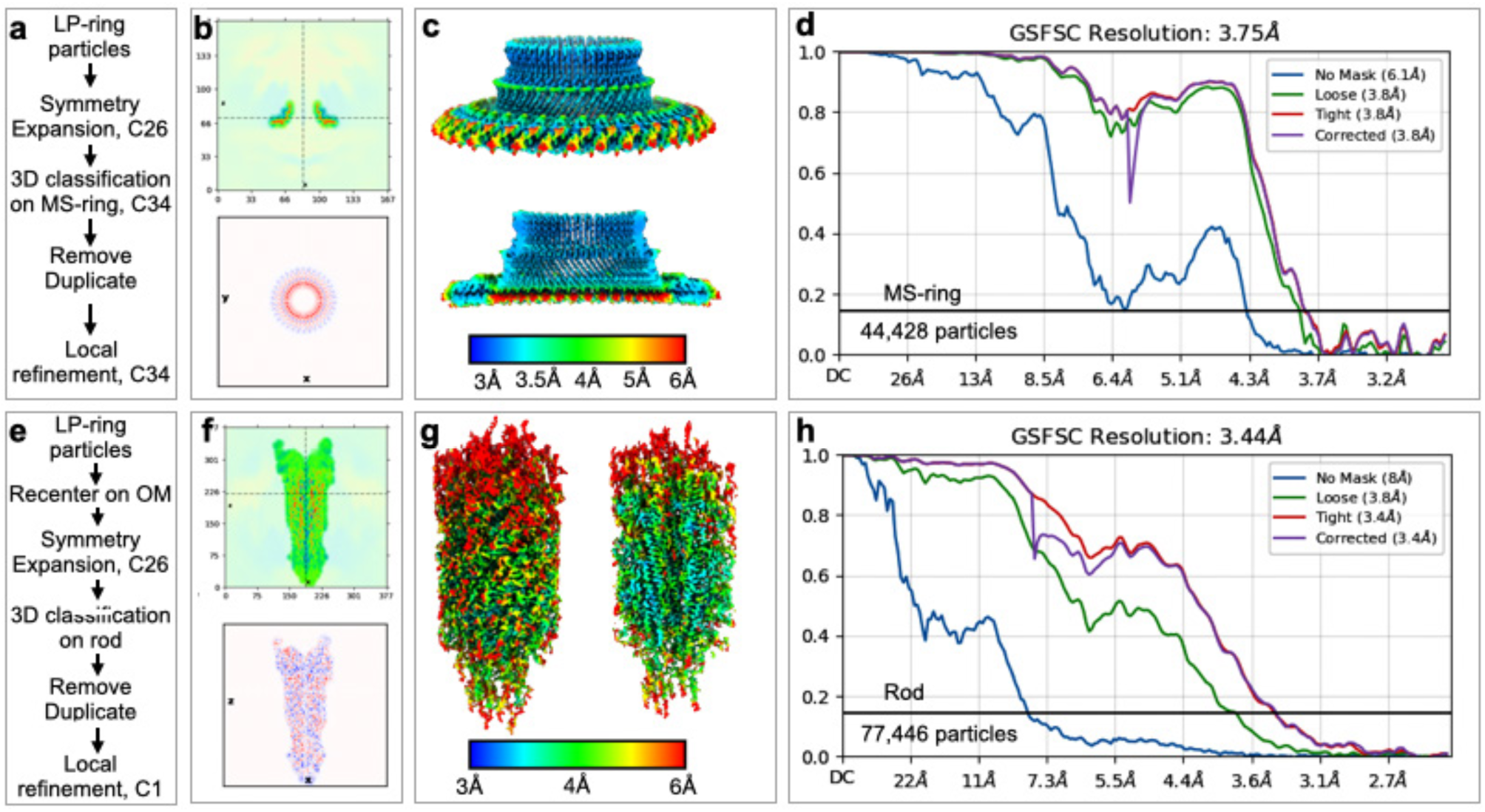
Data processing of the MS-ring and rod. **(a)** Workflow for MS-ring structural determination. **(b-d)** Focused mask and local resolution map of the RBM3 domain of FliF in MS-ring, indicating an overall resolution of 3.75 Å from 44,428 particles. **(e)** Workflow for rod structural determination with C1 symmetry. **(f-h)** Focused mask and local resolution map of the rod, determined from 77,446 particles at C1 symmetry. FSC curve of the rod indicates an overall resolution of 3.44 Å.

**Fig. S4:**
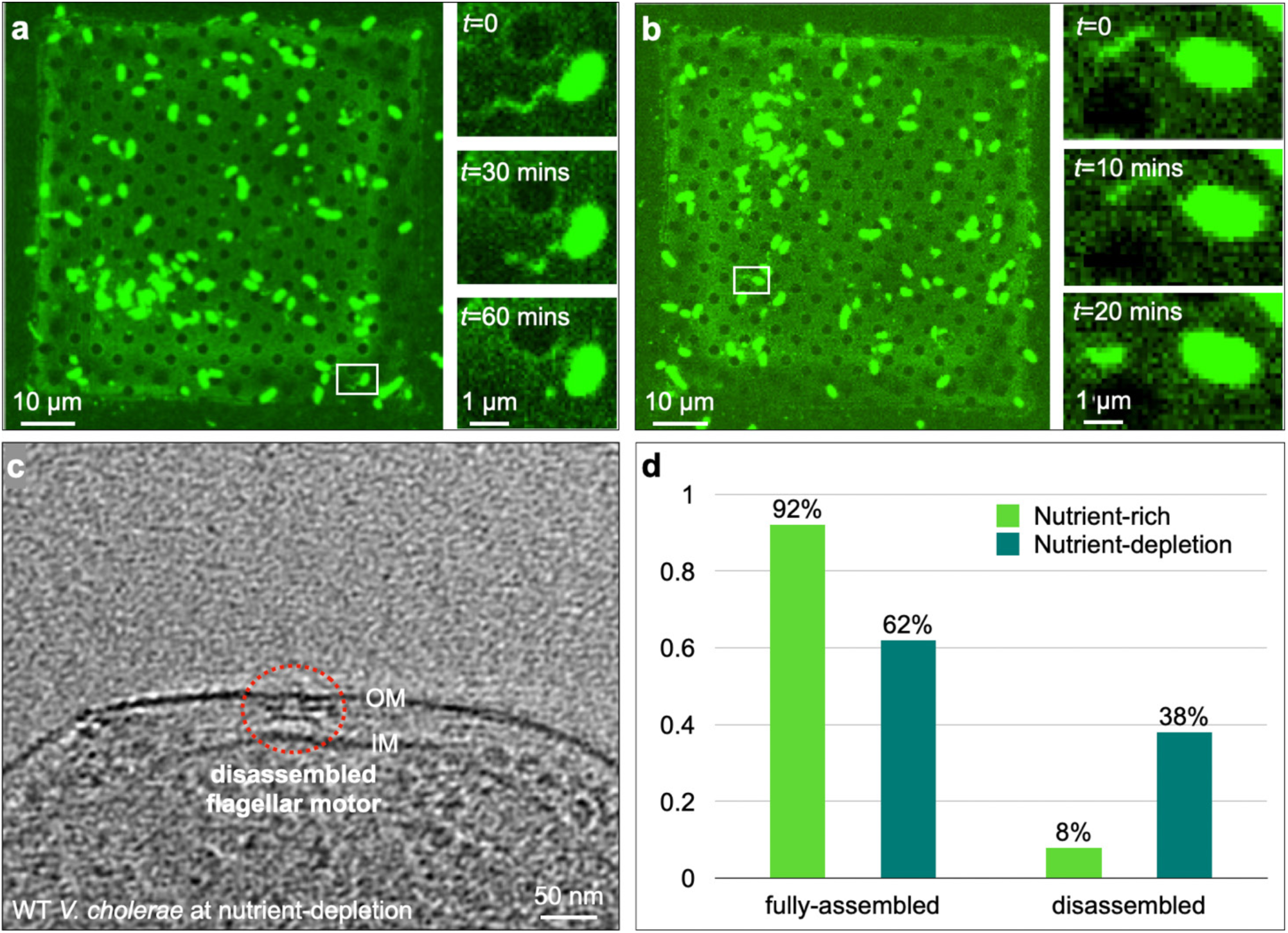
Light microscopy showing flagellar shedding in live cells. **(a-b)** Two reprehensive confocal microscopy imaging of flagellar shedding at 30- and 10-minute intervals. **(c)** Cryo-ET snapshot of a cell tip showing flagellar detachment under the same conditions as panel (**a**, **b**). **(d)** Quantification of flagellar disassembly under nutrient-rich (LB medium) and nutrient-depleted (M9 medium) conditions that trigger flagellar shedding. Experiments were performed in the WT strain. A total of 81 cells were analyzed under nutrient-depletion and 273 cells under nutrient-rich conditions.

**Fig. S5:**
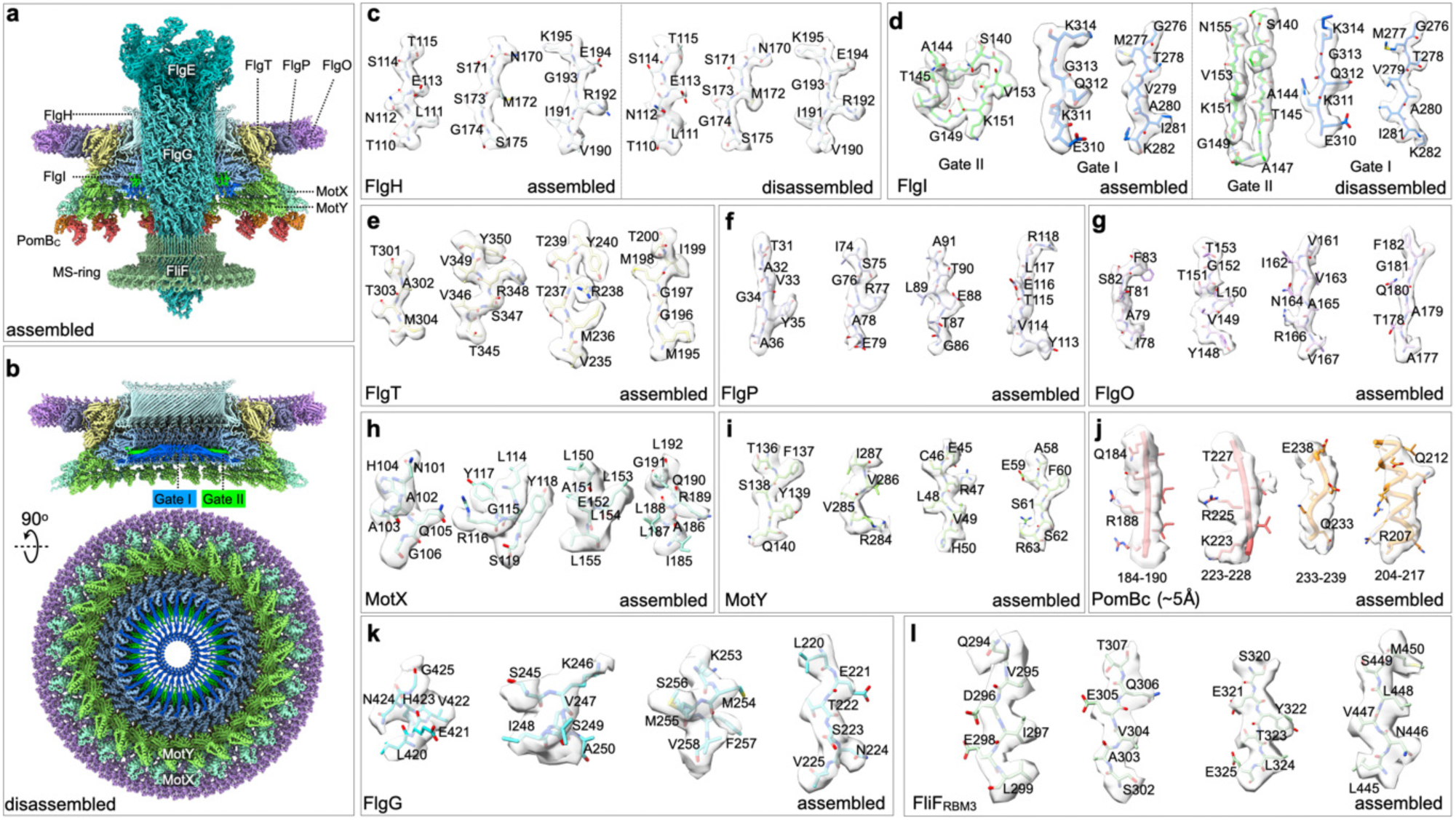
Model-to-map fitting and representative densities at different resolutions. **(a, b)** *de novo* atomic models of the flagellar outer membrane complex both at assembled and disassembled states comprised of 12 flagellar proteins with ∼340 units. Representative atomic models were fitted into cryo-EM densities from the Δ*flhG V. cholerae* strain. Models are shown as sticks superimposed on cryo-EM densities (transparent gray surface), visualized in ChimeraX with the ‘volume zone’ set to a 2 Å cutoff. **(c, d)** Identical model segments of FlgH and FlgI fitted into density maps of the assembled and disassembled flagellar motors, respectively. GateI and GateII regions in FlgI are highlighted. **(e-l)** Model-to-map fits of FlgT, FlgP, FlgO, MotX, MotY, FlgG in the rod, and the RBM3 domain of the FliF in MS-ring for the assembled flagellar motor. The resolution for PomBC is low, only backbone and partial sidechains fitting into PomBC density is shown.

**Fig. S6:**
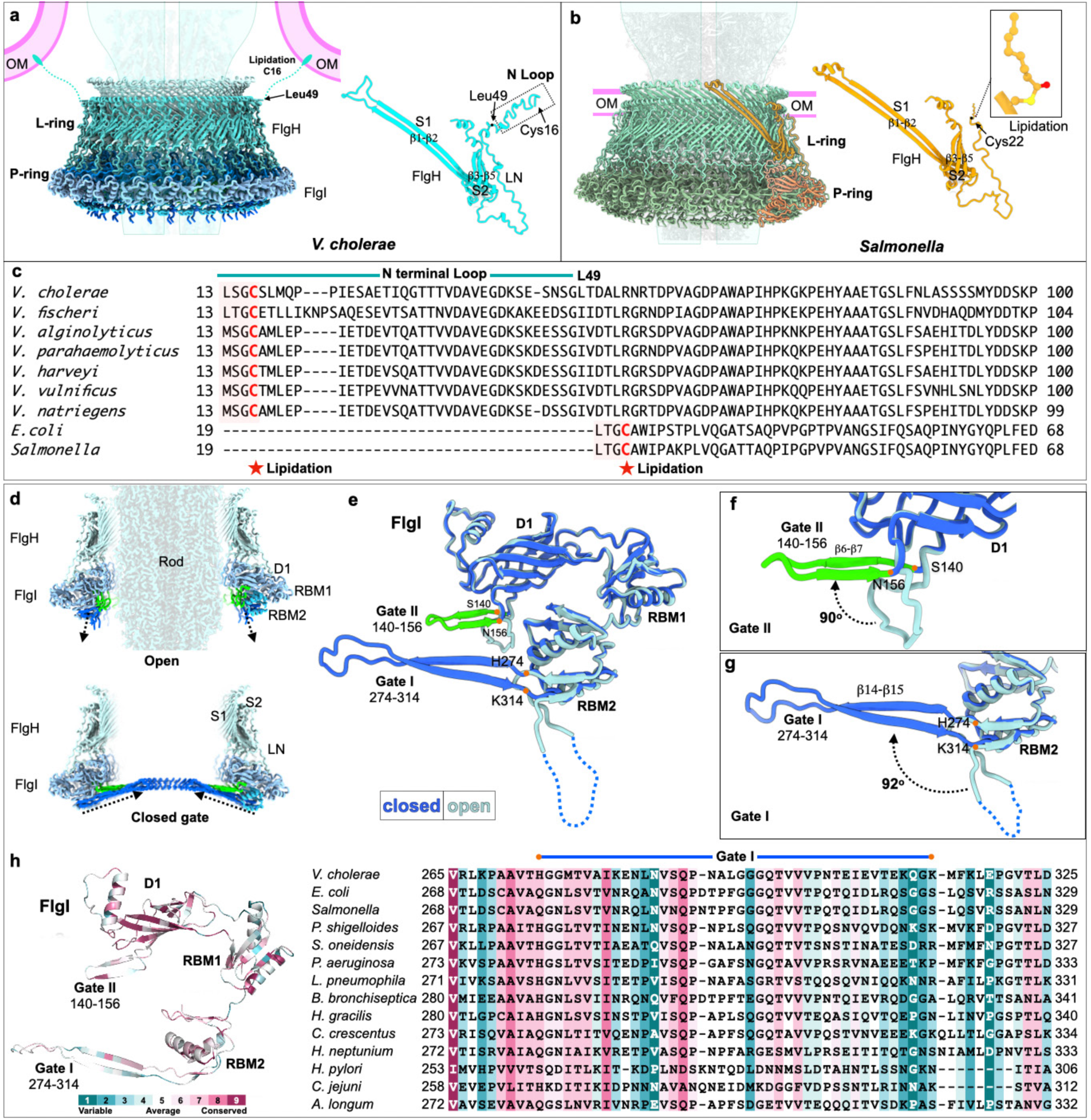
LP-ring model of assembled and disassembled flagellar motors showing open and closed configurations of FlgI. **(a)** LP-ring model of *V. cholerae* at the assembled state. Monomer of FlgH shows the flexible region of C16-L49, where are predicted to form a flexible loop that may extend outward and insert lipid-modified Cys16 into the inner leaflet of the outer membrane. **(b)** LP-ring structures in *S. enterica*. The FlgH is stabilized by the outer membrane through lipidation at Cys22 of FlgH. **(c)** Sequence alignment of FlgH from *Vibrio* species compared with unsheathed flagella from *E. coli* and *S. enterica*. **(d)** Cross-sectional views of the LP-ring in assembled and disassembled states, illustrating the open and closed configurations of FlgI. In the assembled state, GateI and GateII tilt downward and away from the rod, creating an open tunnel that allows the rod to pass through. In the disassembled state, both gates tilt upward toward the rod, forming a closed configuration. **(e)** Zoom-in views of GateI and GateII. In the open state, residues H274-K314 of GateI are disordered and invisible, while in the closed state, they form an antiparallel β-sheet. GateII remains ordered in both states, forming a flexible loop in the open state and a short antiparallel β-sheet in the closed state. **(f, g)** From open to closed state, GateI and GateII tilt upward toward the rod by ∼92° and ∼90°, respectively. **(h)** Conservation of FlgI sequences across different bacterial species.

**Fig. S7:**
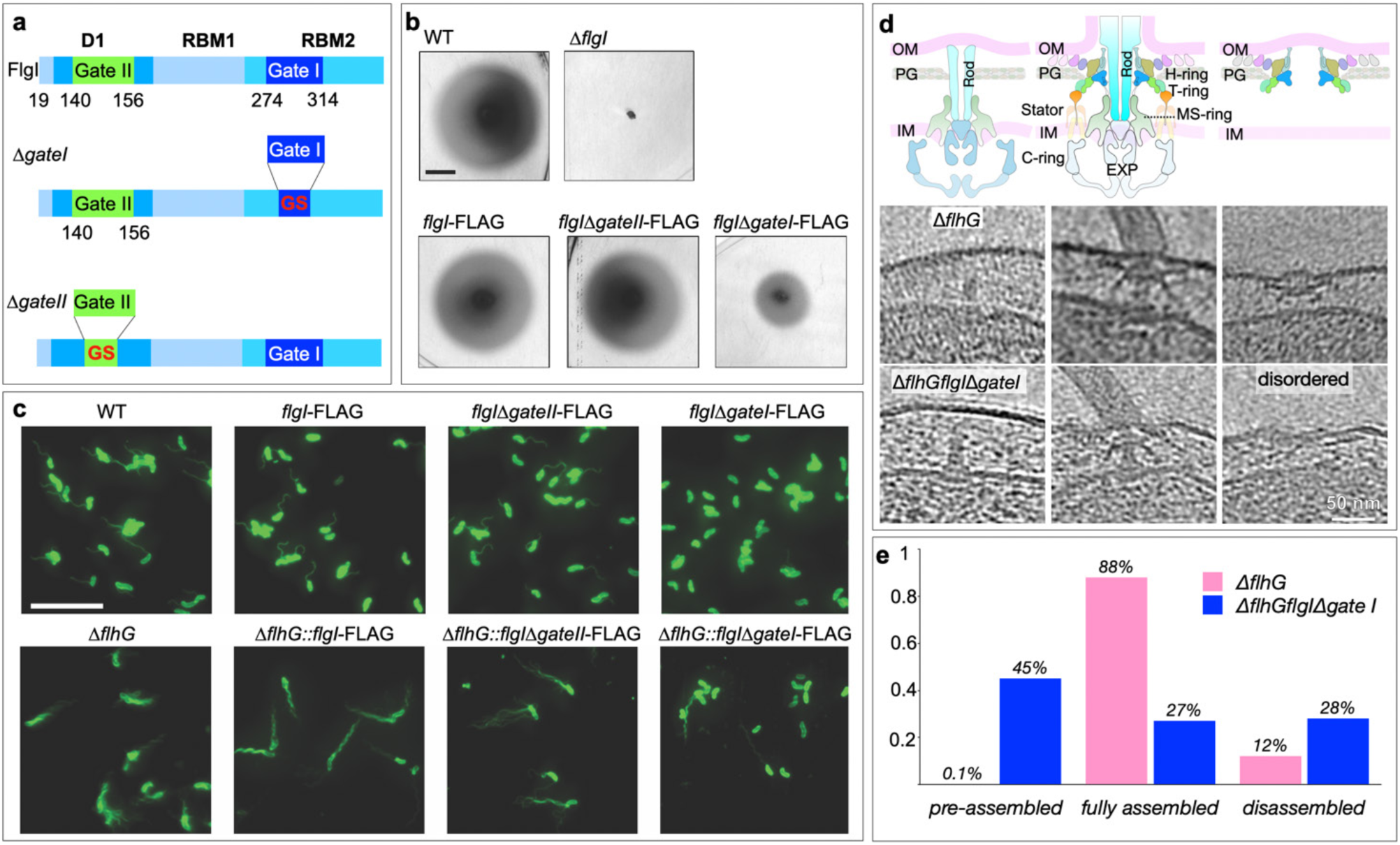
Motility assay and cryo-ET snapshots of *flgI* mutants. **(a)** Mutation of GateI (H274–K314) and GateII (S140–N156) in FlgI to examine their roles in flagellar assembly. *flgI*Δ*gateI* and *flgI*Δ*gateII* represent mutants with GateI and GateII replaced by a glycine-serine (GS) linker, respectively. **(b)** Motility assay of *flgI* mutants on LB soft agar plates. Representative images of migration zones for the mutants are presented. Scale bar = 1 cm. **(c)** Representative fluorescence microscopy images showing flagella stained with a membrane dye in *flgI* variants in WT and Δ*flhG* backgrounds. Scale bar = 10 µm. **(d)** Cartoon model and cryo-ET snapshots illustrating representative assembly stages of the flagellar motor: pre-assembled, fully assembled, and disassembled. The phenotype of Δ*flhGflgI*Δ*gateII* matches that of the Δ*flhG* mutant and is not shown. The pre-assembled state contains only the basal body and rod, without the T-ring, H-ring, or other components. Fully assembled and disassembled states correspond to our two resolved atomic structures. Compared with Δ*flhG*, the disassembled motor in Δ*flhGflgI*Δ*gateI* appears fuzzy, likely due to the absence of GateI. **(e)** Quantification of cells in pre-assembled, fully assembled, and disassembled states for Δ*flhG* (*n* = 151) and Δ*flhG flgI*Δ*gateI* (*n* = 112) mutants.

**Fig. S8:**
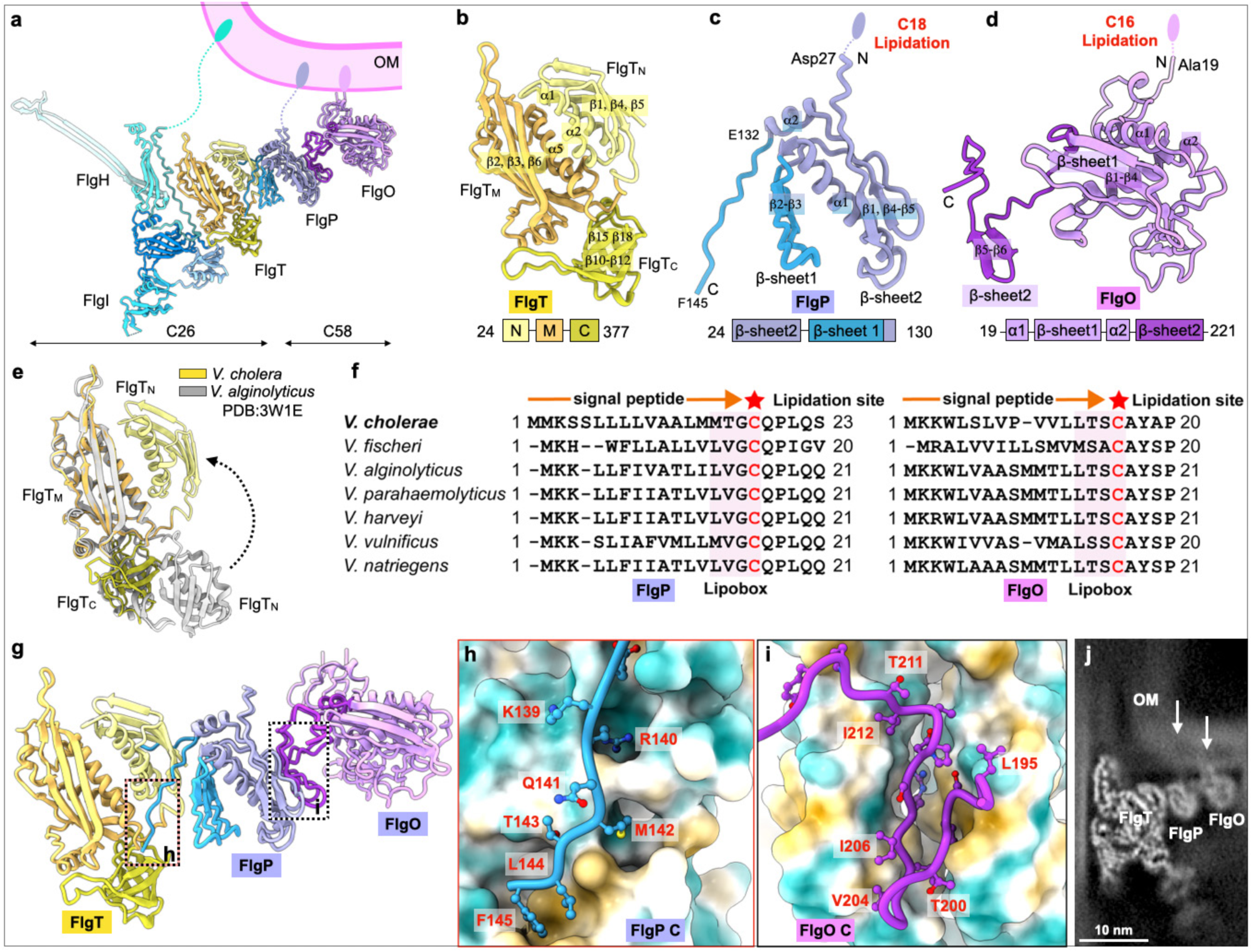
Near-atomic model of H-ring. **(a)** Monomers of the LP-ring and H-ring. FlgH, FlgI, and FlgT exhibit C26 symmetry, whereas FlgP and FlgO exhibit C58 symmetry. **(b)** Atomic model of FlgT (residues 24-377), divided into three domains: FlgTN (A24-T109), FlgTM (K121-C287), and FlgTC (P292-L377). FlgTC lies beneath FlgTM and serves as a scaffold for T-ring assembly. **(c)** Atomic model of FlgP (residues 24-130), consisting of two β-sheets (β2-β3; β1, β4-β5) and two α-helices (α1, α2). The N-terminal Cys18 lipidation is highlighted to be anchored to the outer membrane (OM), and the region from C18 to A27 is too flexible to solve. **(d)** Atomic model of FlgO, containing four β-strands (β1-β4) and two α-helices (α1, α2) in a compact fold. A long N-terminal loop precedes α1 and extends toward the OM till A19. The lipidation Cys16 is labeled to be anchored to the OM. **(e)** Superposition of FlgT models from *V. cholerae* and *V. alginolyticus* (PDB: 3W1E)(*20*). **(f)** Sequence alignment of N-terminal domains of FlgP and FlgO in *Vibrio* species. **(g)** Overview of the interactions among the H-ring subunits. **(h)** Zoom-in view of the C-terminal loop of FlgP interacts with FlgT. **(i)** Zoom-in view of the C-terminal loop of FlgO interacts with FlgP. **(j)** Cryo-EM density map of FlgO and FlgP showing their attachment with OM.

**Fig. S9:**
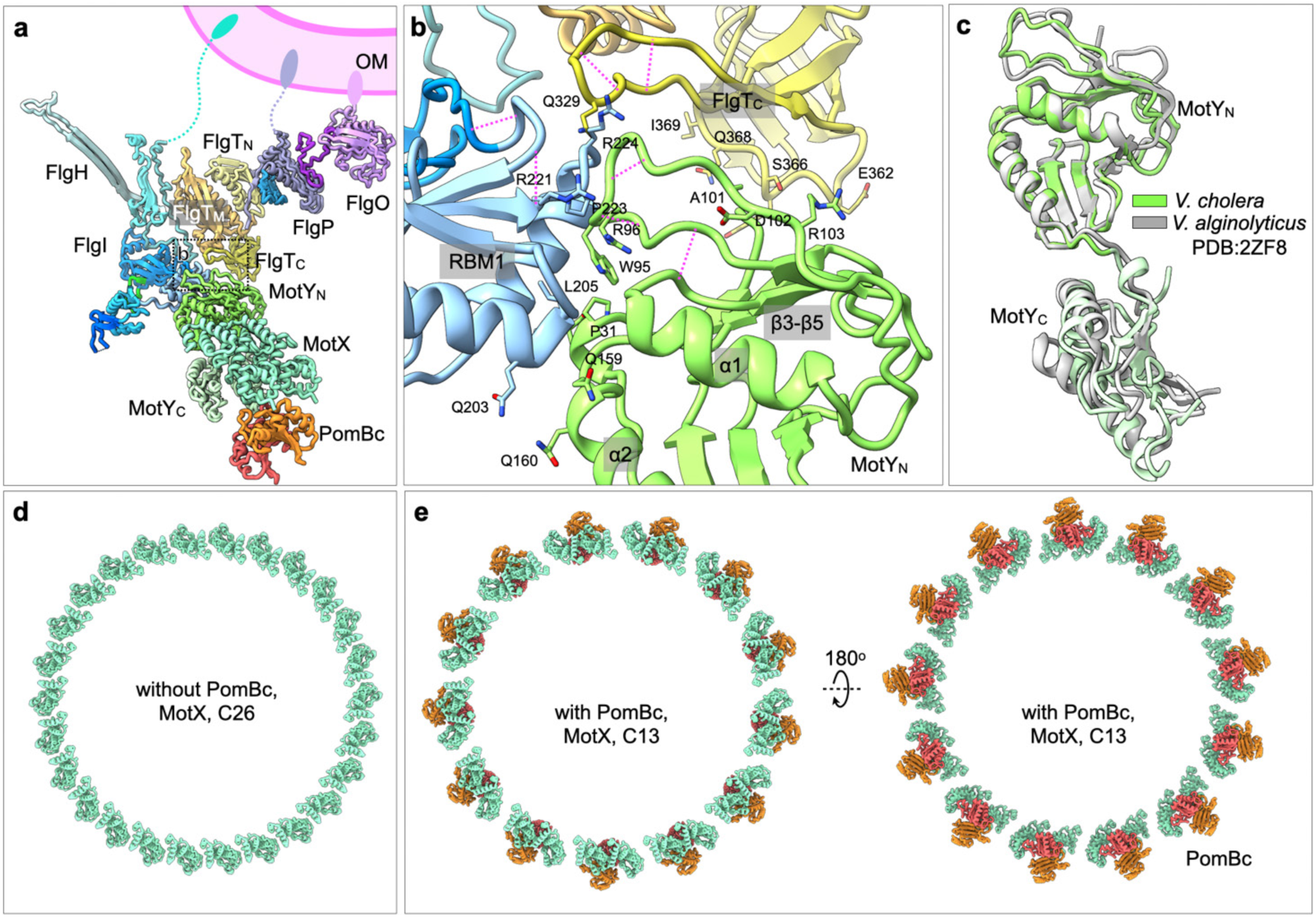
Near-atomic model of T-ring and PomBc. **(a)** Monomer of the T-ring and PomBC located below the LP-ring and H-ring. **(b)** Interaction sites among MotYN, FlgTC, and the RBM1 domain of FlgI. The magenta dashed lines indicate hydrogen bonds. **(c)** Superposition of MotY from *V. cholerae* and *V. alginolyticus* determined by X-ray diffraction (PDB: 2ZF8)(*21*). **(d)** MotX ring with C26 symmetry in the absence of PomBC. **(e)** Top and bottom views of MotX and PomBC in C13 symmetry. Upon engagement with PomBC, two MotX subunits move closer to form a dimer, which together with PomBC forms a bundled complex.

**Fig. S10:**
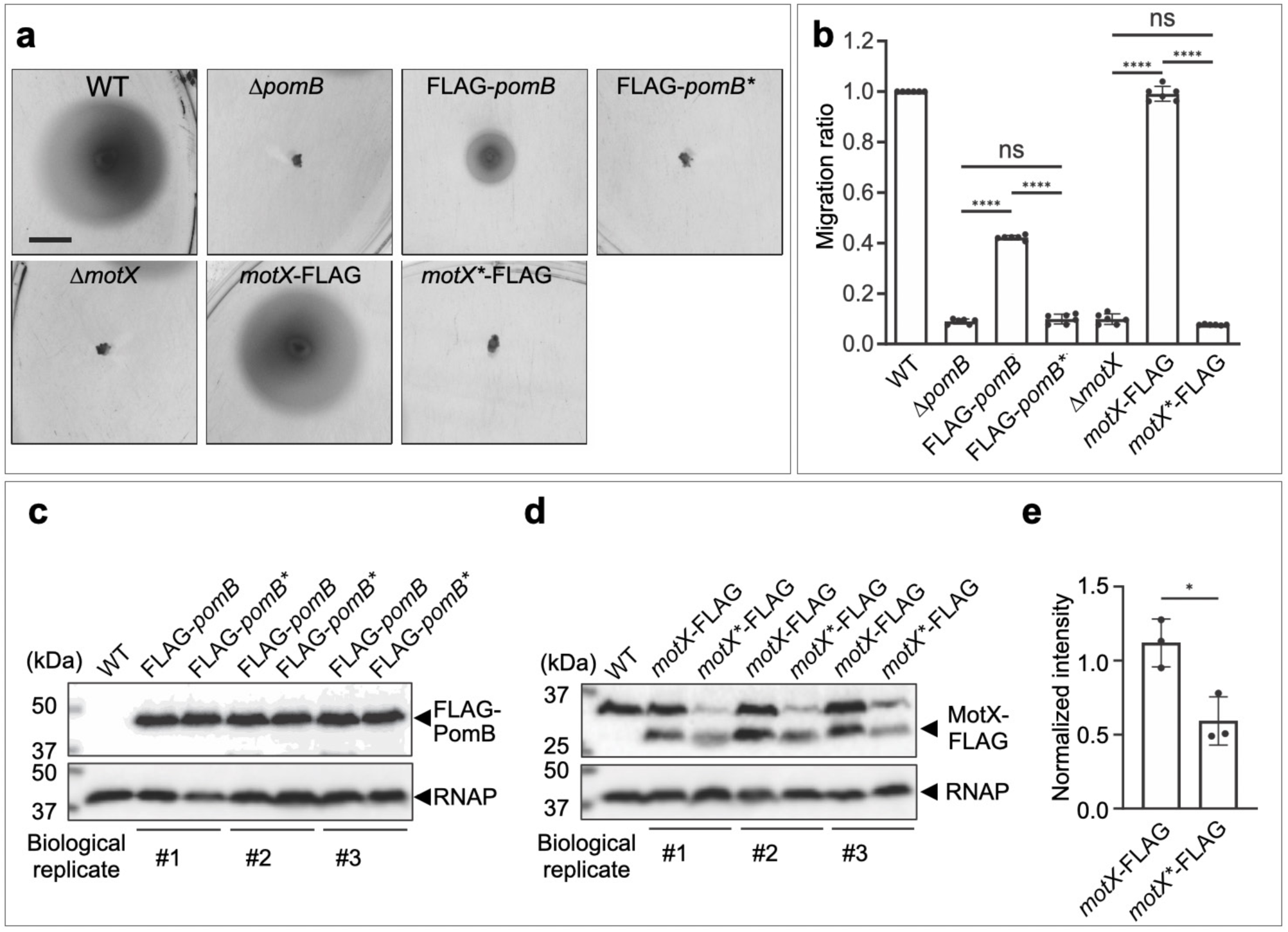
PomB and MotX protein abundance and motility on LB soft agar plates. **(a, b)** Western blot analysis of *pomB* (a) and *motX* (b) mutants. The *pomB* mutant (PomB^P196A,F201A,Q203A,F206A,L239A^) is denoted as *pomB\**, and the *motX* mutant (MotX^P29A,I58A,Q60A,D61A,A64R,I69A^) is denoted as *motX\**. To determine whether these substitutions affect protein abundance, the FLAG epitope was introduced into PomB and MotX and their abundance was assessed by western blot analysis. All *pomB* alleles were tagged with an N-terminal FLAG epitope, while all *motX* alleles were tagged with a C-terminal FLAG epitope. Data are representative of three biological replicates. The expected sizes of each protein are as follows: FLAG-PomB/PomB* (39 kDa), MotX/MotX*-FLAG (28 kDa), and RNAP (36 kDa). These are indicated by arrows in the figure. **(c)** Quantification of the *motX* mutant western blot data shown in panel (b). Band intensities were measured and normalized to the intensity of the loading control (RNAP). Statistical significance was determined using an unpaired t-test. Means from individual biological replicates (*n* = 3) of *motX*\*-FLAG were compared to those of *motX*-FLAG. Adjusted P values ≤ 0.05 were deemed significant. *, *p* ≤ 0.05. FLAG-PomB and FLAG-PomB* are produced at comparable levels, whereas MotX-FLAG* is present at ∼50% of the MotX-FLAG abundance. **(d)** Motility of *pomB* and *motX* mutants on LB soft agar plates. Representative images of migration zones for the mutants are presented. Scale bar = 1 cm. **(e)** Quantification of soft agar migration zones shown in panel (d), presented as migration ratios for the mutants. Statistical significance was determined using a One-Way ANOVA with Tukey’s correction for multiple comparisons. Means from individual biological replicates (*n* = 6) were compared to that of WT. Adjusted P values ≤ 0.05 were deemed significant. ****, *p* ≤ 0.0001; ns, not significant. FLAG**-***pomB* migrates at an intermediate level between WT and Δ*pomB*, indicating that an N-terminal FLAG tag reduces PomB function. By contrast, FLAG-*pomB*\* is essentially non-motile and phenocopies Δ*pomB*, consistent with a non-functional PomB*. *motX*-FLAG migrates similarly to WT, indicating that MotX-FLAG is functional. *motX\**-FLAG is non-motile and phenocopies Δ*motX*.

**Fig. S11:**
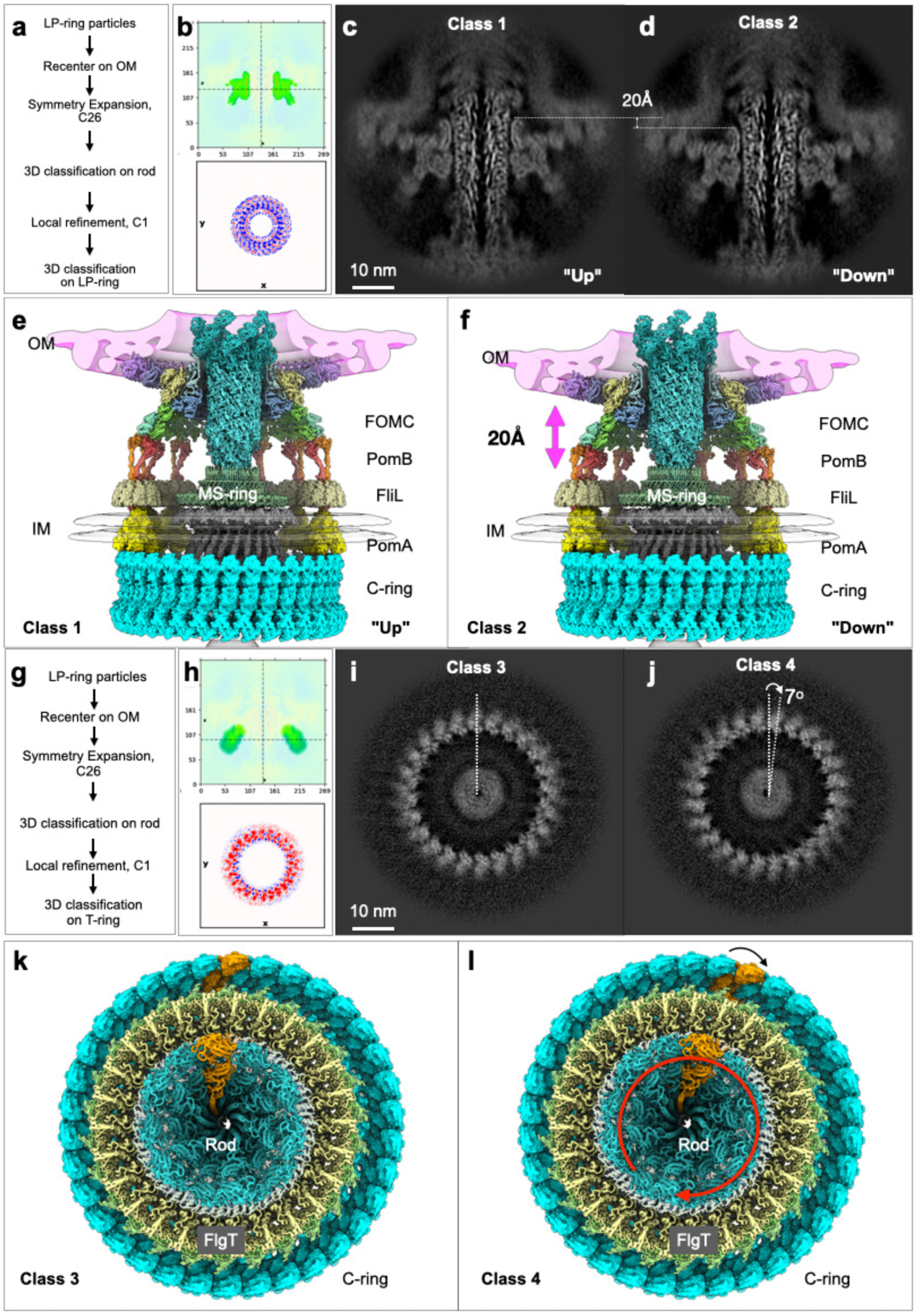
3D classification distinguishes multiple conformations of the flagellar outer membrane complex (FOMC) and rod. **(a)** Workflow of 3D classification on LP-ring, which is based on coordinates of rod to evaluate LP-ring flexibility relative to the rod. **(b)** Focused mask. **(c, d)** Two representative classes of the LP-ring, showing that while the rod remains steady, the LP-ring adopts “up” and “down” conformations resembling a sliding motion on the rod. **(e, f)** Corresponding atomic model of complete flagellar motor. The FliL(*22*), PomA(*23*) and C-ring(*24*) is adapted from published work. The FOMC slides up to ∼20 Å relative to the rod in different rotation stages. **(g)** Workflow of 3D classification on T-ring, which is also based on coordinates of rod. **(h)** Focused mask on the T-ring. **(i, j)** Two representative classes of the T-ring, showing that identical subunits differ in orientation by ∼7°, resembling of a rotating motion of rod relative to the FOMC. **(k, l)** Atomic model of complete flagellar motor at two rotating states, corresponding to panel (i, j). A subunit of FlgE in hook, FliG in C-ring were highlighted in orange to illustrate the rotation of C-ring, rod and hook, while the FOMC remains fixed. FlgO, FlgP, MotX, MotY and PomB in the FOMC were removed to better visualize the rotor components.

**Table S1:**
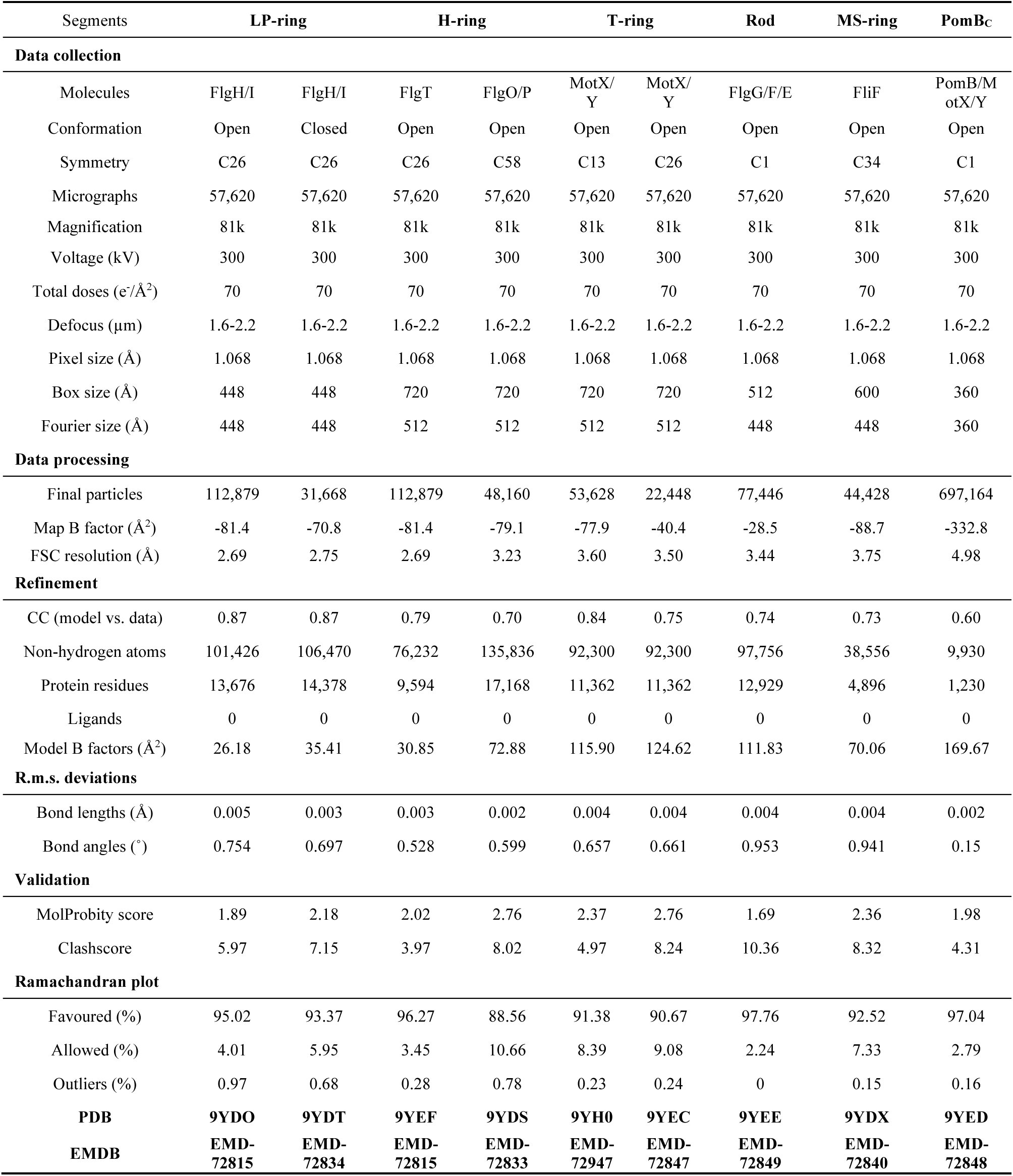
Cryo-EM data collection, processing in cryoSPARC and structural refinement.

| Segments | LP-ring |  | H-ring |  | T-ring |  | Rod | MS-ring | PomBc |
| --- | --- | --- | --- | --- | --- | --- | --- | --- | --- |
| Data collection |  |  |  |  |  |  |  |  |  |
| Molecules | FlgH/I | FlgH/I | FlgT | FlgO/P | MotX/<br>Y | MotX/<br>Y | FlgG/F/E | FliF | PomB/M<br>otX/Y |
| Conformation | Open | Closed | Open | Open | Open | Open | Open | Open | Open |
| Symmetry | C26 | C26 | C26 | C58 | C13 | C26 | C1 | C34 | C1 |
| Micrographs | 57,620 | 57,620 | 57,620 | 57,620 | 57,620 | 57,620 | 57,620 | 57,620 | 57,620 |
| Magnification | 81k | 81k | 81k | 81k | 81k | 81k | 81k | 81k | 81k |
| Voltage (kV) | 300 | 300 | 300 | 300 | 300 | 300 | 300 | 300 | 300 |
| Total doses (e<sup>-</sup>/Å<sup>2</sup>) | 70 | 70 | 70 | 70 | 70 | 70 | 70 | 70 | 70 |
| Defocus (μm) | 1.6-2.2 | 1.6-2.2 | 1.6-2.2 | 1.6-2.2 | 1.6-2.2 | 1.6-2.2 | 1.6-2.2 | 1.6-2.2 | 1.6-2.2 |
| Pixel size (Å) | 1.068 | 1.068 | 1.068 | 1.068 | 1.068 | 1.068 | 1.068 | 1.068 | 1.068 |
| Box size (Å) | 448 | 448 | 720 | 720 | 720 | 720 | 512 | 600 | 360 |
| Fourier size (Å) | 448 | 448 | 512 | 512 | 512 | 512 | 448 | 448 | 360 |
| Data processing |  |  |  |  |  |  |  |  |  |
| Final particles | 112,879 | 31,668 | 112,879 | 48,160 | 53,628 | 22,448 | 77,446 | 44,428 | 697,164 |
| Map B factor (Å<sup>2</sup>) | -81.4 | -70.8 | -81.4 | -79.1 | -77.9 | -40.4 | -28.5 | -88.7 | -332.8 |
| FSC resolution (Å) | 2.69 | 2.75 | 2.69 | 3.23 | 3.60 | 3.50 | 3.44 | 3.75 | 4.98 |
| Refinement |  |  |  |  |  |  |  |  |  |
| CC (model vs. data) | 0.87 | 0.87 | 0.79 | 0.70 | 0.84 | 0.75 | 0.74 | 0.73 | 0.60 |
| Non-hydrogen atoms | 101,426 | 106,470 | 76,232 | 135,836 | 92,300 | 92,300 | 97,756 | 38,556 | 9,930 |
| Protein residues | 13,676 | 14,378 | 9,594 | 17,168 | 11,362 | 11,362 | 12,929 | 4,896 | 1,230 |
| Ligands | 0 | 0 | 0 | 0 | 0 | 0 | 0 | 0 | 0 |
| Model B factors (Å<sup>2</sup>) | 26.18 | 35.41 | 30.85 | 72.88 | 115.90 | 124.62 | 111.83 | 70.06 | 169.67 |
| R.m.s. deviations |  |  |  |  |  |  |  |  |  |
| Bond lengths (Å) | 0.005 | 0.003 | 0.003 | 0.002 | 0.004 | 0.004 | 0.004 | 0.004 | 0.002 |
| Bond angles (°) | 0.754 | 0.697 | 0.528 | 0.599 | 0.657 | 0.661 | 0.953 | 0.941 | 0.15 |
| Validation |  |  |  |  |  |  |  |  |  |
| MolProbity score | 1.89 | 2.18 | 2.02 | 2.76 | 2.37 | 2.76 | 1.69 | 2.36 | 1.98 |
| Clashscore | 5.97 | 7.15 | 3.97 | 8.02 | 4.97 | 8.24 | 10.36 | 8.32 | 4.31 |
| Ramachandran plot |  |  |  |  |  |  |  |  |  |
| Favoured (%) | 95.02 | 93.37 | 96.27 | 88.56 | 91.38 | 90.67 | 97.76 | 92.52 | 97.04 |
| Allowed (%) | 4.01 | 5.95 | 3.45 | 10.66 | 8.39 | 9.08 | 2.24 | 7.33 | 2.79 |
| Outliers (%) | 0.97 | 0.68 | 0.28 | 0.78 | 0.23 | 0.24 | 0 | 0.15 | 0.16 |
| PDB | 9YDO | 9YDT | 9YEF | 9YDS | 9YH0 | 9YEC | 9YEE | 9YDX | 9YED |
| EMDB | EMD-72815 | EMD-72834 | EMD-72815 | EMD-72833 | EMD-72947 | EMD-72847 | EMD-72849 | EMD-72840 | EMD-72848 |

**Table S2:** Bacterial strains and plasmids used in this study.

| Strain or plasmid | Relevant genotype | Antibiotic<sup>R</sup> | Source |
| --- | --- | --- | --- |
| ***E. coli* strains** |  |  |  |
| DH5α λpir | *endA1 hsdR17 glnV44 (=supE44) thi-1 recA1 gyrA96 relA1 ϕ80dlacΔ(lacZ)M15 $\Delta$(lacZYA-argF)U169 *zdg-232::Tn10 uidA::pir*<sup>+</sup>* | | (25) |
| S17-1 λpir | *Tp<sup>r</sup> Sm<sup>r</sup> recA thi pro r<sub>K</sub><sup>-</sup> m<sub>K</sub><sup>+</sup> RP4::2-Tc::MuKm Tn7 λpir* | Tp<sup>R</sup>, Sm<sup>R</sup> | (26) |
| SM10 λpir | *thi thr leu tonA lacY supE recA* (RP4-2-Tc::Mu) λpirR6K Km<sup>r</sup> $\pi$<sup>+</sup> | Km<sup>R</sup> | (27) |
| ***V. cholerae* strains** |  |  |  |
| FY_VC_00001 | *Vibrio cholerae* O1 El Tor A1552, WT | Rif<sup>R</sup> | (28) |
| FY_VC_00337 | Δ*flaA* | Rif<sup>R</sup> | This study |
| FY_VC_18435 | Δ*flhG* | Rif<sup>R</sup> | This study |
| FY_VC_19705 | Δ*flhG*Δ*flgI* | Rif<sup>R</sup> | This study |
| FY_VC_19703 | Δ*flgI* | Rif<sup>R</sup> | This study |
| FY_VC_19691 | Δ*pomB* | Rif<sup>R</sup> | This study |
| FY_VC_19693 | Δ*motX* | Rif<sup>R</sup> | This study |
| FY_VC_19707 | *flgI*-FLAG | Rif<sup>R</sup> | This study |
| FY_VC_19709 | *flgI*<sup>*</sup> (=flgI<sup>G141-I152</sup>→GS)-FLAG | Rif<sup>R</sup> | This study |
| FY_VC_19711 | *flgI*<sup>**</sup> (=flgI<sup>H272-K314</sup>→GS)-FLAG | Rif<sup>R</sup> | This study |
| FY_VC_19713 | Δ*flhG::flgI*-FLAG | Rif<sup>R</sup> | This study |
| FY_VC_19715 | Δ*flhG::flgI*<sup>*</sup>-FLAG | Rif<sup>R</sup> | This study |
| FY_VC_19717 | Δ*flhG::flgI*<sup>**</sup>-FLAG | Rif<sup>R</sup> | This study |
| FY_VC_19695 | FLAG-*pomB* | Rif<sup>R</sup> | This study |
| FY_VC_19697 | FLAG-*pomB*<sup>*</sup> (=pomB<sup>P196A,F201A,Q203A,F206A,L239A</sup>) | Rif<sup>R</sup> | This study |
| FY_VC_19699 | *motX*-FLAG | Rif<sup>R</sup> | This study |
| FY_VC_19701 | *motX*<sup>*</sup> (=motX<sup>P29A,I58A,Q60A,D61A,A64R,I69A</sup>)-FLAG | Rif<sup>R</sup> | This study |
| JY016 | *Vibrio cholerae* O1 biovar El Tor strain C6706 *str2* | Sm<sup>R</sup> | (29) |
| **Plasmids** |  |  |  |
| pGP704*sacB*28 | pGP704 derivative, *mob/oriT sacB* | Amp<sup>R</sup> | (30) |
| pFY_332 | pGP704*sacB*28::Δ*flaA* | Amp<sup>R</sup> | This study |
| pFY_7382 | pGP704*sacB*28::Δ*flhG* | Amp<sup>R</sup> | This study |
| pFY_7875 | pGP704*sacB*28::Δ*flgI* | Amp<sup>R</sup> | This study |
| pFY_5310 | pGP704*sacB*28::Δ*pomB* | Amp<sup>R</sup> | (31) |
| pFY_5883 | pGP704*sacB*28::Δ*motX* | Amp<sup>R</sup> | (31) |
| pFY_7876 | pGP704*sacB*28::*flgI*-FLAG | Amp<sup>R</sup> | This study |
| pFY_7877 | pGP704*sacB*28::*flgI*<sup>*</sup>-FLAG | Amp<sup>R</sup> | This study |
| pFY_7878 | pGP704*sacB*28::*flgI*<sup>**</sup>-FLAG | Amp<sup>R</sup> | This study |
| pFY_7871 | pGP704*sacB*28::FLAG-*pomB* | Amp<sup>R</sup> | This study |
| pFY_7873 | pGP704*sacB*28::FLAG-*pomB*<sup>*</sup> | Amp<sup>R</sup> | This study |
| pFY_7872 | pGP704*sacB*28::*motX*-FLAG | Amp<sup>R</sup> | This study |
| pFY_7874 | pGP704*sacB*28::*motX*<sup>*</sup>-FLAG | Amp<sup>R</sup> | This study |

**Table S3:**
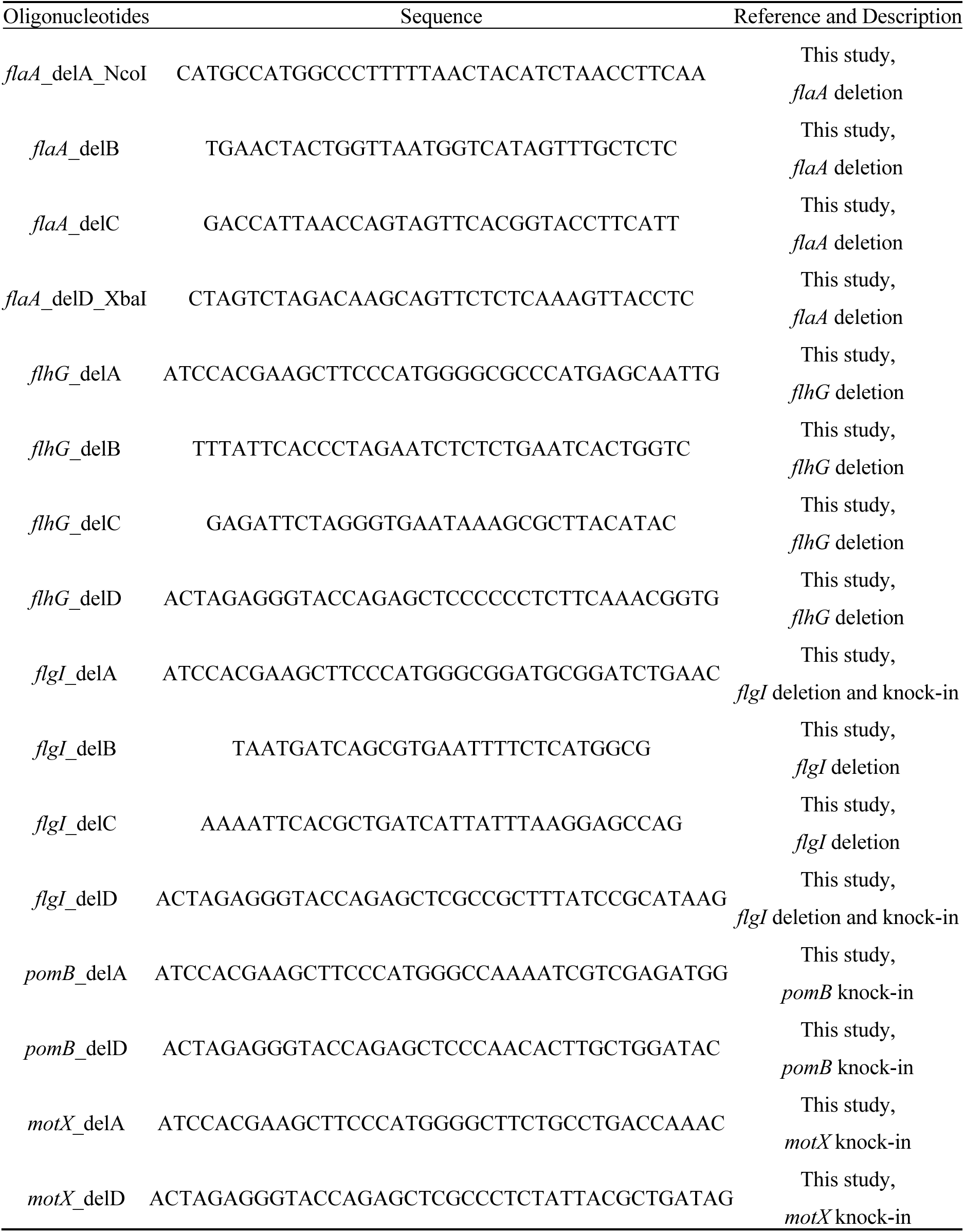
Oligonucleotides used in this study.

**Table S4:** Cryo-ET data collection, processing.

| **Bacterial strains** | **WT** | **WT** | **$\Delta flhG$** | **$\Delta flhG flgI \Delta gateI$** | **$\Delta flhG flgI \Delta gateII$** |
| --- | --- | --- | --- | --- | --- |
| Culture condition | LB | M9 | LB | LB | LB |
| Magnification | 42k | 42k | 42k | 42k | 42k |
| Voltage (kV) | 300 | 300 | 300 | 300 | 300 |
| Total doses ($e^-/\text{\AA}^2$) | 70 | 70 | 70 | 70 | 70 |
| Defocus ($\mu\text{m}$) | -4.8 | -4.8 | -4.8 | -4.8 | -4.8 |
| Tomograms | 273 | 81 | 442 | 112 | 84 |

**Movie S1: Animation of assembly of the sheathed flagellum.** Subtomogram averages in left panel provide evidence that, as the hook elongates, it gradually pushes the outer membrane outward, resulting in the formation of a continuous sheath that envelops the growing hook. The emerging hook extends perpendicularly from the outer membrane in a straight conformation.

**Movie S2: Animation of dynamic rotation and ejection of the sheathed flagellum.**

## Notes

### Competing Interest Statement

The authors have declared no competing interest.

